# Divergent selection on dispersal targets chemosensory and neuronal genes in *Tribolium castaneum*

**DOI:** 10.1101/2025.08.17.670711

**Authors:** Michael D. Pointer, Will J. Nash, Lewis G. Spurgin, Mark McMullan, Simon Butler, David S. Richardson

## Abstract

Dispersal is key to the life history, ecology and evolution of many organisms, and important in pest invasiveness. However, the genetic architecture underlying variation in dispersal behaviour remains poorly understood outside of a few model species. We investigated the genomic basis of dispersal using artificial selection on replicated lines of *Tribolium castaneum*, a flour beetle, an emergent model system, and an economically important agricultural pest. Combining whole-genome resequencing with population-level genotype-phenotype association analysis, we identify genomic regions associated with selection on dispersal. Identified candidate genes were significantly enriched for functions related to neuronal structure and function, as well as chemosensory behaviour and mating, suggesting that variation in dispersal is mediated by neural and chemosensory pathways. Our results demonstrate that dispersal propensity has a polygenic basis and support an interaction between dispersal and mating ecology in this system. These findings contribute to a deeper understanding of the genetic mechanisms driving dispersal evolution of dispersal and its role in shaping eco-evolutionary dynamics.

## Introduction

Dispersal is a complex life history trait with a critical role in the ecology and evolution of many species (Ronce 2007). Individual movements influence population size, density, and range (Kokko and López-Sepulcre 2006), and thus the metapopulation dynamics that contribute to population persistence or expansion in fragmented landscapes (Clobert 2012; Legrand *et al*. 2017). Furthermore, via effects gene flow, dispersal determines patterns of genetic variation within and among populations, with implications for their evolutionary trajectories (Holsinger and Weir 2009). Dispersal is also an important aspect of evolutionary responses to anthropogenic climate change, habitat fragmentation, the dynamics of biological invasions and agricultural pest distributions (Travis *et al*. 2013; Legrand *et al*. 2017; Renault *et al*. 2018). Thus, knowledge of the genetic underpinnings of dispersal is essential to understand the causes and consequences of dispersal evolution (Ronce 2007).

Across taxa, dispersal is commonly seen as a component of suites of coevolving phenotypes, referred to as behavioural syndromes (Clobert 2012). Individual variation in such syndromes may represent different life-history strategies (Sih *et al*. 2004). Patterns of association between these traits are complex, highly context-dependent, and the mechanistic basis of such associations is poorly understood (Clobert 2012). Despite this, recent studies have revealed that, under conditions such as range expansion, dispersal itself can rapidly evolve over short timescales (Ochocki and Miller 2017; Weiss-Lehman *et al*. 2017; Simcox *et al*. 2024). For example, dispersal evolution during the cane toad (*Bufo marinus*) invasion of Australia has accelerated the advance of the range front by fivefold in less than 100 years (Shine *et al*. 2021). Uncovering the genetic architecture of dispersal traits will clarify how variation underlying dispersal is maintained in populations, aid in elucidating the basis of behavioural syndromes, inform our ability to trace how molecular variation leads to phenotypic differences in movement patterns, and enable predictions around the adaptive potential of dispersal (Saastamoinen *et al*. 2018).

The genetic architecture of traits is key to how they respond to selection (Pritchard and Di Rienzo 2010; Le Corre and Kremer 2012). The genetic basis of dispersal-related traits has been described in a range of species, revealing highly varying genetic architectures (Saastamoinen *et al*. 2018; Dochtermann *et al*. 2019). Dispersal is usually thought to be a polygenic trait (Merilä and Sheldon 1999; Pritchard and Di Rienzo 2010), an idea supported by work in both insects and vertebrates (Jordan *et al*. 2012; Saatoglu *et al*. 2024). For example, ∼300 genes were associated with dispersal phenotype in Mountain Pine Beetles (*Dendroctonus ponderosae;* Shegelski et al. 2021). However, in some taxa large-effect loci influence dispersal (Saastamoinen *et al*. 2018), which can be separated into those with metabolic (Niitepõld and Saastamoinen 2017) or neurophysiological (Sokolowski 1980; Trefilov *et al*. 2000; Fidler *et al*. 2007; Krackow and König 2008; Anreiter and Sokolowski 2019) effects on movement. Notable examples include the *For* gene (*Foraging*; and its homologues), a neuro-signalling regulator linked to movement behaviour in taxa from *C.elegans* to humans (reviewed in Anreiter and Sokolowski 2019); and *Pgi*, which underlies phenotypic variation in flight metabolism and dispersal propensity in wild butterflies (reviewed in Niitepõld and Saastamoinen 2017). Simulation studies modelling the rate of dispersal evolution under different architectures have also provided conflicting results (Saastamoinen *et al*. 2018; Weiss-Lehman and Shaw 2022).

Replicated experimental evolution, in combination with resequencing (Schlötterer *et al*. 2015), and powerful statistical methods enables genotype-phenotype associations to be resolved (Coop *et al*. 2010; Gautier 2015; Olazcuaga *et al*. 2020). The emergent genomic model *Tribolium castaneum* is highly suited to such experimental evolution studies (Pointer *et al*. 2021; Campbell *et al*. 2022). The species is a globally significant pest, responsible for large economic losses and impacts on food security (Phillips and Throne 2010), consequently its dispersal ecology is of great applied interest. Previous work on *T. castaneum* has indicated that dispersal may be a component of a behavioural syndrome, covarying with key life-history traits (Lavie and Ritte 1978; Zirkle *et al*. 1988; Pointer *et al*. 2024). In this species, individual-level dispersal variation seems to be driven by activity levels and movement patterns (Pointer, Spurgin, Vasudeva, *et al*. 2024), suggesting that, as in other systems, the phenotype may stem from differences in neurophysiology and/or metabolism (Saastamoinen *et al*. 2018). Some genes whose expression covaries with walking motivation have been identified in *T. castaneum* (Matsumura *et al*. 2024), but as of yet no study has investigated the molecular genomic basis of dispersal.

Here, we utilise whole genome re-sequencing of individuals from replicated lines of *T. castaneum* (n=32), previously selected for divergent dispersal propensity, and displaying robustly repeatable behaviour (Pointer *et al*. 2023; Pointer *et al*. 2024), to identify the genetic architecture of adaptation to selection on dispersal. We employ BayPass, leveraging the power of the study’s population-level replication, to highlight candidate SNPs significantly associated with dispersal selection regimes. Signatures of suppressed nucleotide diversity around a subset of identified candidates support the recent occurrence of selective sweeps in these regions. We then functionally enrich sets of candidate genes to explore how molecular variation might be linked to dispersal phenotypes. We find many genes associated with dispersal phenotype, supporting a polygenic trait architecture. The functions of the candidate genes suggest that neuronal structure and function affecting chemosensation are the principal mechanisms underpinning dispersal evolution in our experiment.

## Methods

### Beetles and husbandry

The Krakow super-strain (KSS) of *Tribolium castaneum* flour beetles was created by combining 14 laboratory populations from across the world, producing a highly outbred strain, the ideal substrate for selection to act upon (Laskowski *et al*. 2015). This strain has been maintained at a census size of 600 individuals for ∼150 generations. All beetle populations were kept on a fodder medium of 90% organic flour 10% brewers yeast, on a constant 12:12 light:dark cycle, with relative humidity of 60% and a regime of non-overlapping generations. At 12±3 days post eclosion, adults are sieved from the fodder and a randomly chosen subset is combined in fresh fodder to begin a seven-day mating and oviposition period, after which adults are removed. Eggs remaining in the fodder then develop over a 35-day development period. By preventing any interaction between sexually mature adults and offspring, this method reduces the risk of negative density-dependent effects, removes the opportunity for intergenerational interactions, such as egg cannibalism, and allows accurate tracking of passing generations. This study complies with applicable UK legislation on sampling from natural populations and animal experimentation (SI 2012/3039).

### Artificial selection for dispersal

Thirty-two experimental lines were founded from KSS stock and artificially selected for dispersal propensity as described by Pointer et al. (2023). Briefly, high (n=16) and low (n=16) dispersal lines were bred under divergent artificial selection over five generations, using a dispersal assay in which each individual, housed within groups of 200, was given three opportunities to ‘disperse’: i.e. leave a patch of suitable habitat, cross a short distance of unsuitable habitat and not return. Individuals that ‘dispersed’ three times out of the three opportunities were considered to display a dispersive phenotype. Individuals that ‘dispersed’ zero times were considered to display a non-dispersive phenotype. Thirty individuals of each of these phenotypes were selected to found the subsequent generation of the relevant treatments, while individuals of intermediate phenotype were discarded.

After a single generation of selection, the mean dispersal phenotype (mean number of dispersals per individual out of three opportunities) between the treatments was significantly different. After five generations of selection, the distribution of the dispersal phenotype between the two treatments was non-overlapping (Pointer *et al*. 2023). Between generations six and 16, a less stringent selection regime was applied. In odd-numbered generations, a reduced assay that allowed a single dispersal opportunity was used to phenotype the dispersers and non-dispersers, of which 30 of the relevant type were selected to found the subsequent generation of each line. In odd-numbered generations, 100 randomly chosen individuals founded the next generation. In generation 17, dispersal was quantified and found to be still strongly divergent and non-overlapping between treatments (Pointer *et al*. 2024; dispersals per individual out of a maximum of three, low dispersal lines = 0.70±0.06; high dispersal lines = 2.44±0.04).

### Sample preparation and sequencing

In generation 17, samples were collected for sequencing from dispersal selection lines and from the ancestral KSS population. Adult females were sampled and flash-frozen in liquid nitrogen. For each individual, DNA extraction was conducted using the DNeasy blood and tissue kit (insect tissue protocol, Qiagen), with the whole individual ground in liquid nitrogen. The extract was then purified using a 1x AMPure XP SPRI (Beckman Coulter) bead cleanup. Library preparation and sequencing were performed at the Earlham Institute (Norwich, UK) using the low-input transposase-enabled pipeline (LITE; see supplementary methods). Libraries were sequenced on two S4 flowcells over two lanes on the Illumina Novaseq 6000 platform. Sequences were obtained from 210 individuals, six from each of 32 dispersal lines and 18 from the KSS stock.

### Variant calling and filtering

Reads were trimmed using Trimmomatic v.039 (Bolger *et al*. 2014) and mapped to the T.cast5.2 reference genome (<u>10.1186/s12864-019-6394-6</u>), using BWA-MEM v0.7.17 (Li 2013). Mapping was followed by SAMtools v1.18 fixmate and SAMtools sort (Danecek *et al*. 2021). PCR duplicates were then removed using Picard v2.26.2 RemoveDuplicates (Broad Institute, 2019). Finally, mappings were filtered for complete read pairs and those with a mapping quality (MAPQ) >25 using SAMtools view.

Joint genotyping was conducted using BCFtools v1.18.0 mpileup (Danecek *et al*. 2021). BCFtools call was then used to call all sites under the multi-allelic model (-m). BCFtools filter was used to remove variants within 3bp of other variants, with a variant quality score <30, that were at a locus with sequencing depth less than 578 and greater than 5201 (+/-3 times total sequencing depth), and were represented by data at that locus in less than 50% of individuals (-g 3 -G 3 -e ’DP < 578 || DP > 5201 || F_MISSING > 0.5 || QUAL < 30’). The resulting file is referred to hereafter as the allsites vcf. From the allsites vcf, single nucleotide polymorphisms (SNPs) were extracted using BCFtools view, and further filtered to remove sites with minor allele count <3. This file is referred to hereafter as the SNP vcf.

To prepare the allsites vcf for further analysis, variant and invariant sites were handled separately: Invariant sites were extracted using VCFtools 0.16.0 (Danecek *et al*. 2011), and stored in a separate file. Variant sites were filtered using VCFtools to only contain biallelic SNPs that did not deviate significantly from Hardy-Weinberg Equilibrium (HWE p-value < 0.001). Following this, invariant and variant sites were concatenated and indexed using BCFtools concat and index. This file is referred to hereafter as the filtered allsites vcf.

QC of the SNP vcf revealed three individual samples (10HT, 11HT3, 11HT4) exhibited huge counts of singleton SNPs and indels compared to other samples. As this is likely a sign of sequencing error, we excluded reads from these samples from the raw sequence data and reran the above steps to regenerate both the SNP vcf and the allsites vcf.

### Linkage disequilibrium

The SNP vcf was down-sampled to 1 SNP per 0.5kb and used to calculate pairwise linkage disequilibrium (LD) between SNPs up to a maximum distance of 5Mb, using VCFtools. As per-line sample sizes were not sufficient to reliably estimate LD, estimates were obtained from ‘populations’ consisting of all individuals from within each treatment (high dispersal, low dispersal, KSS) and within all treatments combined. Using a custom R script (see data availability for link to the Github repository, we calculated mean LD within distance bins of 1kb and plotted this to visualise LD in the data (figure S1). In each treatment LD halved from the maximum at ∼50kb, we therefore took this as a window size with which to begin to examine patterns across the genome.

### Population structure

As a measure of artefact detection and replicate verification, principal component analysis (PCA) was performed using plink (v1.9; (Purcell *et al*. 2007). The SNP vcf was pruned for linkage with bcftools (+prune -m 0.3 -w 50kb) before PCA was conducted with plink2 (Chang *et al*. 2015).

### Identification of candidate loci

We performed a genome-wide scan for selection using BayPass v2.4 (Gautier 2015), implementing a Bayesian framework sensitive to demography. BayPass estimates a background allele frequency (omega) matrix across populations to account for the confounding effect of demography, which can frustrate the identification of selected variants (Günther and Coop 2013; Gautier 2015). This approach allowed us to control for unquantified differences in relatedness among individuals used to found each selection line. The BayPass model uses an omega matrix, calculated from neutral SNPs, to correct for population covariation when testing allele frequencies for population divergence or association with environmental or trait variables. Within BayPass, we utilised the statistic, C_2_, which contrasts allele frequencies between two groups of populations specified by a binary trait (Olazcuaga *et al*. 2020). This method outperforms others in identifying SNPs under selection (Olazcuaga *et al*. 2020). We computed C_2_ across our 32 dispersal lines, with the dispersal selection treatment as the binary covariable.

To avoid the impact of small, annotation-sparse, unplaced scaffolds in the reference genome, we ran BayPass on the 10 linkage-group-level Tcas5.2 scaffolds. We computed the BayPass omega matrix using a curated subset of 12,232 independent, high confidence, non-exonic SNPs, at putatively neutral loci across the genome, affording the best opportunity to estimate the neutral covariance in allele frequencies (omega dataset; see supplementary methods). The foreground dataset used for BayPass analysis contained a more permissive set of 3,240,899 SNPs, derived from the SNP vcf. This set leveraged BayPass’s robustness to missing data whilst maximising the number of high-confidence SNPs in the analysis (see supplementary methods).

We performed two independent Baypass runs with different random seed initiators and computed correlations to test the consistency of model performance with our data <u>(Dickson et al. 2020; Olazcuaga et al. 2020)</u>. The C_2_ estimates were calibrated using a pseudo-observed dataset (POD; Gautier 2015; see supplementary methods). The 0.999 quantile of C_2_ values from the POD analysis was used as the outlier threshold for empirical C_2_ values. C_2_ candidate SNPs were those with C_2_ above this threshold, and C_2_ candidate regions were defined as those containing >=2 outlier SNPs separated by <50kb (Gautier 2015).

Nucleotide diversity (π) in each replicate population was computed in 10kb non-overlapping windows along the genome using *pixy*, from the allsites VCF. Mean π per window was calculated across the 16 individual populations within each treatment. To identify regions of low π, we computed the mean across all windows for each linkage-group level scaffold; outlier windows were those with mean π more than four standard deviations below the scaffold mean. Trends were visualised using ggplot in R v4.3.3 (see data availability for link to the Github repository), with rolling mean π calculated over 5 windows.

We thereby generated two sets of candidate genes, those within 1) C_2_ candidate regions identified by BayPass, and 2) a subset of C_2_ candidate regions that overlapped a π outlier window.

### Characterisation of candidate genes

Genes associated with selection candidates were identified by intersecting their positions with the annotation (<u>Tribolium_castaneum.T.cas5.2.59.gff3</u>; bedtools *intersect*). The two sets of candidate genes (1 and 2 described above) were used as input. Returned gene lists were used as input to g:Profiler (Reimand *et al*. 2007; Kolberg *et al*. 2023) to test for enrichment of functional terms derived from gene ontology (GO). All other settings were the G:profiler defaults and the background used was all genes in the Tcas5.2.59 annotation. The OSG3 annotation (https://ibeetle-base.uni-goettingen.de/download/species/Tcas/OGS3.gff.gz) and the iBeetle-Base database (Dönitz *et al*. 2018) were used to manually identify gene functions, and orthologous genes in *Drosophila* were identified using Flybase (Öztürk-Çolak *et al*. 2024).

## Results

### DNA sequencing

Following adapter trimming, 199,248 - 32,028,177 reads per sample were mapped to the Tcas5.2 reference. Following removal of PCR duplicates and quality filtering 106,340 - 13,399,500 reads per sample remained, representing 0.09 - 11.83x mean coverage (table S1). The filtered allsites vcf contained 37,840,898 sites, the SNP vcf contained 4,418,680 SNPs.

### Linkage disequilibrium

Linkage disequilibrium (LD) was similar in high dispersal, low dispersal and all treatments combined, peaking at Ca. R^2^=0.2, and halving from the maximum at 0.6-0.7Mb (figure S1A;B;C). The KSS control samples had the highest LD, with a peak of 0.25 and halving at ∼1.3Mb (figure S1D), although this estimate is less reliable due to smaller sample size (n=18 vs. n>=93).

### Population structure

An LD pruned and MAF filtered set of 223,034 SNPs was used to perform principal component analysis. PCA showed no unexpected batch effects (figure S2).

### Identification of candidate loci

BayPass analysis showed high repeatability, with the C_2_ estimates associated with SNPs being highly correlated across two replicate runs with different starting seeds (Pearson’s r = 0.96). Of the 3,240,899 sites in the dataset, we identified 267 SNPs as C_2_ outliers, with representation across all ten linkage groups (figure 1A). Linkage groups two, three, and four contained the most outlier SNPs (44, 93, and 32, respectively), with 88 of those on LG3 being within a single peak. Grouping SNPs following Galthier (2015), we recovered 22 candidate regions associated with dispersal, representing 256 candidate SNPs (figure S5). Nine of the 22 candidate regions overlapped a π outlier window (figure 1, table S2), with the strongest signals of suppressed nucleotide diversity associated with BayPass outlier regions on LG3 (figure 1Bi) and LG7 (figure 1Bii).

**Figure 1.**
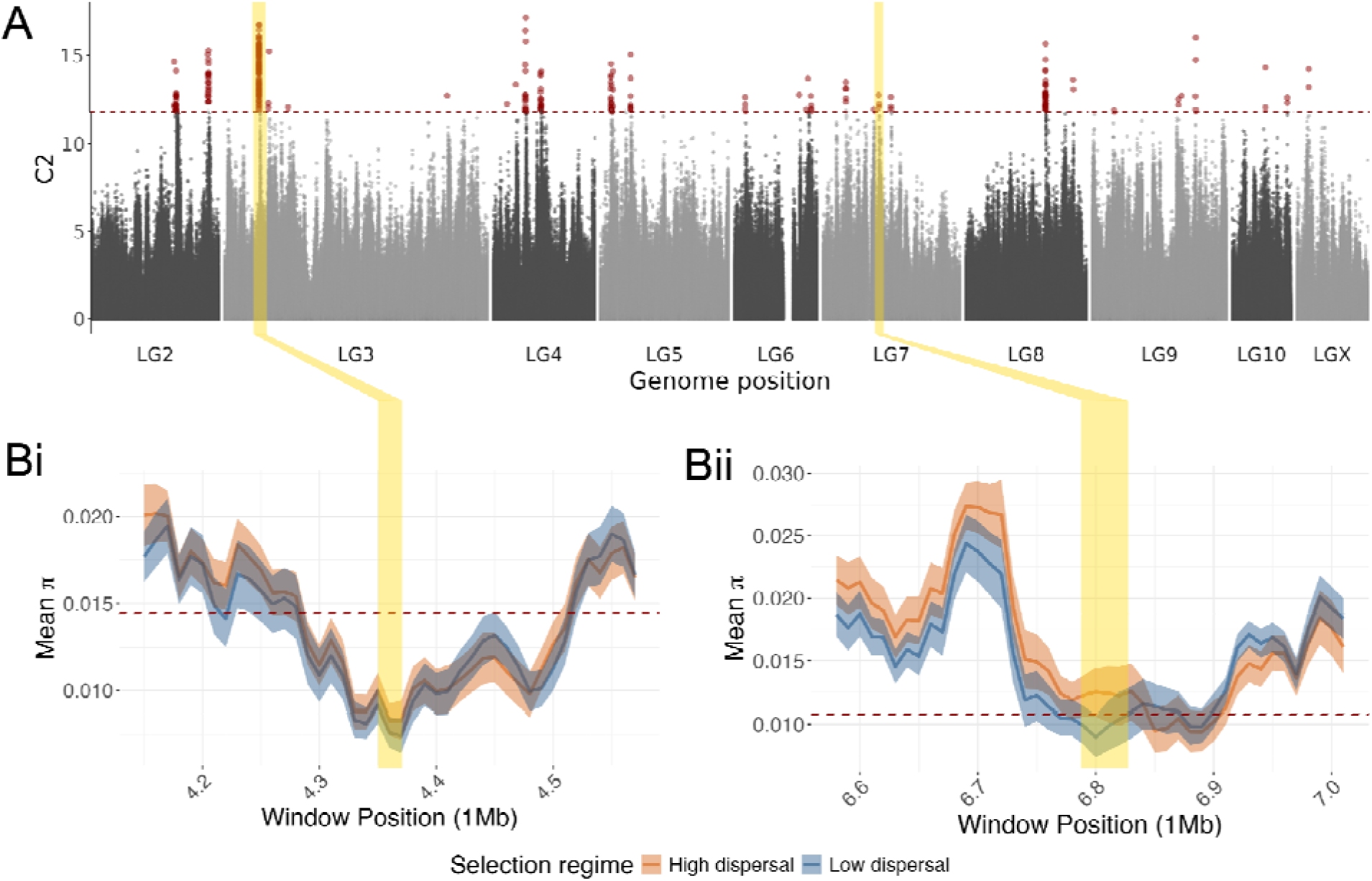
Genomic variation associated with divergent artificial selection for dispersal propensity in 32 independently evolving lines of *Tribolium castaneum*. A) Values of the C_2_ statistic generated by BayPass analysis as a measure of SNP associations with the direction of selection. Red points are in excess of the 0.999 quantile of C_2_ values from a BayPass run on a pseudo-observed dataset of putatively neutral SNPs, shown by the dotted line. **B)** Expanded view of regions on linkage groups three (i) and seven (ii), where BayPass peaks coincide with regions of low nucleotide diversity (π), computed in 10kb windows within each population and averaged across populations from the same selection regime. Dotted red lines represent the mean π of both selection regimes combined on the focal chromosome minus four standard deviations (0.01% of windows are expected to fall beneath this threshold). Yellow shading links the region of interest across panels and indicates the extent of the BayPass outlier regions in B.

### Characterisation of candidate genes

GO analysis was performed independently on two sets of genes. Genes with the strongest evidence of responding to dispersal selection (supported by C_2_ and π) showed enrichment for functions related to neurons, chemosensory behaviour and mating (table 1). The broader set (supported by C_2_ alone) was enriched for functions related to protease activity, wounding response, reproduction, transport and DNA processing.

**Table 1.** Gene ontology analysis using G:profiler to identify functional enrichment among genes in candidate regions located by BayPass and nucleotide diversity analyses, between two sets of *Tribolium castaneum* populations artificially selected for divergent dispersal propensity.

| Gene set | GO/KEGG category | GO/KEGG term | P<sub>adj</sub> |
| --- | --- | --- | --- |
| Genes in C<sub>2</sub> candidate regions overlapping $\pi$ outlier windows (n = 42) | GO:MF | Primary active transmembrane transporter activity | <0.01 |
|  | GO:BP | Reproductive process | <0.05 |
|  | GO:BP | Flagellated sperm motility | <0.05 |
|  | GO:BP | Sperm motility | <0.05 |
|  | GO:BP | Courtship behaviour | <0.05 |
|  | GO:BP | Chemosensory behaviour | <0.05 |
|  | GO:BP | Male courtship behaviour | <0.05 |
|  | GO:BP | Reproductive behaviour | <0.05 |
|  | GO:BP | Male mating behaviour | <0.05 |
|  | GO:BP | Mating behaviour | <0.05 |
|  | GO:CC | Neuronal cell body | <0.05 |
|  | GO:CC | Cell body | <0.05 |
|  | GO:CC | Dendrite | <0.05 |
|  | GO:CC | Dendritic tree | <0.05 |
|  | GO:CC | Somatodendritic compartment | <0.05 |
|  | GO:CC | Axon | <0.05 |
| Genes in C<sub>2</sub> candidate regions (n = 104) | GO:MF | Serine-type endopeptidase activity | <0.001 |
|  | GO:MF | Serine-type peptidase activity | <0.001 |
|  | GO:MF | Serine hydrolase activity | <0.001 |
|  | GO:MF | Endopeptidase activity | <0.001 |
|  | GO:MF | Peptidase activity | <0.001 |
|  | GO:MF | DNA topoisomerase type I (single strand cut, ATP-independent) activity | <0.05 |
|  | GO:BP | Blood coagulation | <0.01 |
|  | GO:BP | Hemostasis | <0.01 |
|  | GO:BP | Coagulation | <0.01 |
|  | GO:BP | Reproduction | <0.01 |
|  | GO:BP | Wound healing | <0.01 |
|  | GO:BP | Regulation of body fluid levels | <0.01 |
|  | GO:BP | Multicellular organism reproduction | <0.05 |
|  | GO:BP | Response to wounding | <0.05 |
|  | KEGG | ABC transporters | <0.05 |

## Discussion

Here, we use replicated lines of red flour beetles, artificially selected for differential dispersal behaviour, to undertake the first study of the genomic basis of this trait in *Tribolium*. Using whole genome resequencing with population-level genotype-phenotype association (GPA) analysis we identify signatures of selection at many regions across the genome, indicating that adaptation was polygenic. Genes within candidate regions were enriched for functions associated with neuronal function and chemosensation, suggesting a possible mechanism underlying dispersal variation.

Previous work in *T. castaneum* suggests that dispersal behaviour may have a relatively simple genetic basis (Pointer *et al*. 2023), possibly a single locus of large effect (Ogden 1970a; Ritte and Lavie 1977). Challenging this suggestion, here we find nine regions across the genome strongly associated with dispersal phenotype, with weaker support for a further 13 regions, indicating that dispersal behaviour is likely a polygenic trait. While a rapid response to selection, as seen in our selection lines, was suggested by Ritte and Lavie (1977) to be an indicator of a simple genetic basis, recent theory indicates that rapid phenotypic change can also result from relatively small shifts across many loci, especially under a strong novel selection pressure (Pritchard and Di Rienzo 2010; Jain and Stephan 2017). Our result aligns with others from insect and vertebrate systems, showing a complex genetic basis of dispersal-related traits. For example, 192 genes were found to be associated with *Drosophila* locomotion (Jordan *et al*. 2012) and ∼300 genes were differentially expressed between dispersal phenotypes of Mountain Pine Beetles (*Dendroctonus ponderosae) (*Shegelski et al. 2021). Similarly, dispersal in the House sparrow (*Passer domesticus*) is polygenic, with a complex basis involving gene x environment interactions (Saatoglu *et al*. 2024).

The regions of the genome with strongest links to dispersal from our analyses were characterised as involved in neuron structure and functioning, and affecting chemosensory behaviour, courtship and reproduction. Dispersal in *Tribolium* is well known to be influenced by the conspecific environment and chemical signals associated with population density (King and Dawson 1972; Pointer *et al*. 2021), with beetles dispersing more readily from high density populations (Ziegler 1978), and from environments with chemical signals of high density, even in the absence of other beetles (Ogden 1970b). Hence, it seems possible that increased sensitivity to chemical cues could lead to greater dispersal propensity for a given population density. This suggests a potential mechanistic link between the candidate genes and dispersal variation, via altered perception or processing of environmental cues, however confirming this would require considerable functional validation.

Given that chemosenation is also key to finding mates in *Tribolium* and across insect systems (Krieger and Breer 1999; Fedina and Lewis 2008), it follows that changes in chemosensation may also affect mating behaviour. Interestingly, a previous study examining reproductive behaviour in these same experimental lines indicated altered male investment in different reproductive strategies with dispersal phenotype, favouring either increased duration or increased frequency of mating (Pointer *et al*. 2024). In the present study, we identify divergence in genes related to sperm motility, supporting a link between dispersal strategy post-copulatory sexual selection. In particular, the gene TC033673 is a homolog of Drosophila’s *Lost Boys* which encodes a flagellar protein determining the likelihood of the ejaculate reaching the female’s sperm storage receptacle (Yang *et al*. 2011).

In addition, the association between genes involved in neural structure and dispersal phenotypes is intriguing, as key traits within dispersal and broader behavioural syndromes are thought to covary via shared neural mechanisms (Sih *et al*. 2004). While further investigation is needed to make robust mechanistic conclusions, it is already known that the focal beetle lines differ in traits such as boldness, and movement pattern (Pointer *et al*. 2024).

To conclude, we recover candidate loci across the genome showing associations with dispersal. Enrichment for functions related to neuron structure and function affecting chemosensation suggests these as likely mechanisms underpinning dispersal variation. In addition, we show that reproductive traits are also responding to dispersal selection, potentially via shared pathways and/or ecological interactions. Our findings highlight how selection acts on dispersal in this system, a representative of the most species-rich order of organisms and an economically important pest. These findings add to our understanding of the evolution of dispersal, a trait at the heart of many key issues in contemporary biology.

## Author Contributions

**Michael D Pointer:** Investigation (lead); Methodology (lead); Formal analysis (lead); Software (lead); Writing – original draft (lead); Writing – review and editing (equal). **Will J Nash:** Investigation (supporting); Methodology (supporting); Software (supporting); Writing – review and editing (equal). **Lewis G Spurgin:** Conceptualization (supporting); Funding acquisition (equal); Supervision (supporting). **Mark McMullan:** Conceptualization (supporting); Supervision (supporting). **Simon Butler:** Project administration (supporting); Supervision (supporting). **David S Richardson:** Supervision (lead); Project administration (lead); Funding acquisition (equal); Writing – review and editing (equal).

## Acknowledgements

This work was funded by a Biotechnology and Biological Sciences Research Council (BBSRC) studentship to MDP (BB/M011216/1). We are grateful to past and present members of the Tribolium lab at the University of East Anglia, without whose dedicated efforts the study of long-term selection lines would not be possible. Sequencing was performed at the Earlham Institute, provided via the Core Capability Grant BB/CCG2220/1 and its constituent work packages (BBS/E/T/000PR9818 and BBS/E/T/000PR9819), and the Core Capability Grant BB/CCG1720/1 and the National Capability at the Earlham Institute BBS/E/T/000PR9816 (NC1—Supporting EI’s ISPs and the UK Community with Genomics and Single Cell Analysis), BBS/E/T/000PR9811 (NC4—Enabling and Advancing Life Scientists in data-driven research through Advanced Genomics and Computational Training), and BBS/E/T/000PR9814 (NC 3 - Development and deployment of versatile digital platforms for ‘omics-based data sharing and analysis). Authors also acknowledge support from BBSRC Core Capability Grant BB/CCG1720/1 and the work delivered via the Scientific Computing group, as well as support for the physical HPC infrastructure and data centre delivered via the NBI Computing infrastructure for Science (CiS) group. Part of this work was delivered via the BBSRC funded National Bioscience Research Infrastructure (BBS/E/ER/23NB0006) at Earlham Institute by members of the Technical Genomics and Core Bioinformatics Groups.

## Data accessibility and benefit-sharing

Sequence data for this project are archived under European Nucleotide Archive (PRJEB90247). Scripts are available on github (https://github.com/mdpointer/Tribolium_dispersal_genomics), and on Dryad (link TBC during submission). This research provides benefits via the sharing of data and results on public databases as described above.

## Notes

### Competing Interest Statement

The authors have declared no competing interest.

