## Supplementary information for "Divergent selection on dispersal targets chemosensory and neuronal genes in *Tribolium castaneum*"

**Supplementary methods**

**Library preparation, performed at the Earlham institute**

A total of 1ng of DNA was combined with 0.9 µl of Tagment DNA buffer, 0.1 µl Tagment enzyme TDE1 and 2 µl nuclease free water in a reaction volume of 5 µl and incubated for 10 minutes at 55˚C. Following the initial incubation, 5 µl of custom barcoded P5 and P7 compatible primers (2µM), 4 µl 5x Kapa Robust 2G reaction buffer B, 0.4 µl 10mM dNTPs, 0.1 µl Kapa Robust 2G enzyme (Sigma Aldrich: KK5005) and 5.5 µl water were mixed, giving a total PCR volume of 20 µl. The DNA was enriched with 14 cycles of PCR (72˚C for 3 minutes, 98°C for 3 minutes, 14 cycles of: 95°C for 10 seconds, 62°C for 30 seconds, 72°C for 3 minutes, final hold at 4 ˚C). Post PCR, the DNA was cleaned up with 1.25x volume of KAPA Pure Beads from Roche (07983298001) utilising the Tecan Fluent 780 liquid handling platform and final libraries were eluted in EB. The size distribution of each library was determined using the Perkin Elmer GX Touch DNA High Sensitivity assay (DNA High Sensitivity Reagent Kit CLS760672), and a smear analysis on a 450-650bp size range was performed, to equimolar pool the libraries. The pool of libraries were then subjected to size selection on a Blue Pippin 1.5% agarose cassette (R2 marker) from SAGE Science (BDF1510) recovering library molecules between 450-650bp. The final pool was quantified by q-PCR and sequenced on one lane of a 300 cycle Illumina NovaSeq 6000 S4 Reagent Kit v1.5 (Illumina 20028312). For this run the library was diluted down to 0.5 nM using EB (10mM Tris pH8.0) in a volume of 30ul before spiking in 1% Illumina phiX Control v3. This was denatured by adding 7ul 0.2N NaOH and incubating at room temperature for 8 mins, after which it was neutralised by adding 8ul 400mM tris pH 8.0. A master mix of DPX1, DPX2, and DPX3 from an Xp 4-lane kit v1.5 (Illumina 20043131) was made and 105ul added to the denatured pool leaving 150ul at a concentration of 100 pM 130ul of which was loaded onto a NovaSeq S4 flow cell using the NovaSeq Xp Flow Cell Dock. The flow cell was then loaded onto the NovaSeq 6000 along with an NovaSeq 6000 S4 cluster cartridge, buffer cartridge, and 300 cycle SBS cartridge. The NovaSeq had NVCS v1.7.5 and RTA v3.4.4 and was set up to sequence 150bp PE reads. The data was demultiplexed and converted to fastq using bcl2fastq2.

**BayPass**

The omega dataset, used to define the background allele frequency matrix, contained a subset of putatively neutral, high-confidence, highly representative, independent SNPs. Putatively neutral sites were obtained by filtering to those lying more than 10kb from exonic regions. Exon locations were extracted from the Ensembl annotation ([Tribolium_castaneum.Tcas5.2.59.gff3.gz](https://ftp.ensemblgenomes.ebi.ac.uk/pub/metazoa/release-59/gff3/tribolium_castaneum/Tribolium_castaneum.Tcas5.2.59.gff3.gz)) and any SNP within these or 10kb in both directions (bedtools slop -b 10000; (Quinlan and Hall 2010) were excluded from the SNP vcf (bcftools -R). Next, sites were filtered to those that remained variant once KSS control samples were removed. We then filtered to sites on the linkage-group-level scaffolds, before removing rare variants (bcftools -- view MAF<0.04). BayPass is robust to missing data, but not when computing the allele frequency matrix (Gautier 2015; Ahrens *et al.* 2018), which requires low missingness and independent loci (Lotterhos 2019). Therefore, to ensure SNPs were representative of every population, we removed any sites missing in >2 individuals in any single population (vcftools –max-missing-count 2). To increase independence, variants were stringently filtered for LD, informed by our linkage estimation (bcftools +prune -m 0.3 -w 50kb).

The dataset used in BayPass analysis runs consisted of the SNP vcf, filtered to sites that were variant in dispersal selection treatments, on linkage-group-level scaffolds and filtered for MAF, but no additional missingness filter or LD prune was applied.

Once filtered omega and analysis datasets were finalised in VCF format, the allele frequencies at these sites were converted to the population allele count format required by BayPass in a custom shell script (using vcftools –counts2).

This allele count file of the omega dataset was then provided to a BayPass run (--gfile), using default settings, to generate the background matrix. The generated matrix (--omegafile) was then fed back to BayPass in a separate run, along with the allele count file of the analysis dataset (--gfile), and the treatment of each population (-contrastfile) to obtain the C_2_ contrast statistic for each SNP. For these analytical runs, SNPs were filtered with the same minor allele frequency and linkage prune parameters as above, leaving 184,842 sites.

The obtained C_2_ estimates were calibrated using pseudo-observed datasets. We used the *simulate.baypass* function from BayPass_utils.R; <https://github.com/andbeck/BayPass/blob/master/baypass_utils.R>) to simulate allele counts for a set of 100,000 SNPs sampled from the allele frequency correlation distribution defined by the omega matrix, computed from our empirical populations. We ran BayPass on this simulated data, supplying the same accompanying files and parameters as for empirical runs. The 0.999 quantile of C2 estimates derived from this ‘neutral simulation’ was then used to define an outlier threshold for the empirical BayPass output.

**Table S1.** Details of sequencing per individual, following the removal of duplicates (Picard v2.26.2 RemoveDuplicates; Broad Institute, 2019) and quality filtering (MAPQ >25 SAMtools view)

| dupes_removed | | | dupes_removed_filtered | | |
| --- | --- | --- | --- | --- | --- |
| number of reads | primary mappings | mean coverage | number of reads | primary mappings | mean coverage |
| 12,213,204 | 11,684,170 | 10.16 | 9,040,962 | 8,995,909 | 7.94 |
| 11,181,508 | 10,728,659 | 9.32 | 8,276,169 | 8,236,634 | 7.26 |
| 10,425,395 | 9,977,670 | 8.69 | 7,919,440 | 7,881,689 | 6.96 |
| 12,104,960 | 11,537,101 | 10.03 | 9,134,284 | 9,089,122 | 8.02 |
| 13,771,468 | 13,180,772 | 11.46 | 10,277,445 | 10,224,109 | 9.03 |
| 10,120,377 | 9,734,410 | 8.51 | 7,291,604 | 7,258,510 | 6.45 |
| 10,362,134 | 9,913,406 | 8.56 | 7,858,414 | 7,819,982 | 6.87 |
| 10,611,604 | 10,166,845 | 8.79 | 7,852,622 | 7,813,241 | 6.87 |
| 9,826,285 | 9,417,418 | 8.16 | 7,405,197 | 7,368,977 | 6.47 |
| 13,005,436 | 12,421,980 | 10.80 | 9,564,395 | 9,511,943 | 8.41 |
| 13,212,085 | 12,640,549 | 10.96 | 9,750,140 | 9,697,072 | 8.57 |
| 14,565,958 | 13,807,107 | 12.06 | 10,954,229 | 10,892,994 | 9.65 |
| 11,890,264 | 11,398,020 | 9.92 | 8,955,488 | 8,910,523 | 7.88 |
| 11,518,764 | 11,025,793 | 9.57 | 8,778,140 | 8,735,137 | 7.69 |
| 14,091,513 | 13,419,350 | 11.63 | 10,281,156 | 10,222,152 | 9.00 |
| 14,237,868 | 13,573,229 | 11.81 | 10,472,325 | 10,411,939 | 9.20 |
| 16,489,244 | 15,826,742 | 13.74 | 12,727,019 | 12,678,179 | 11.13 |
| 14,365,689 | 13,787,213 | 12.07 | 10,924,327 | 10,884,351 | 9.67 |
| 16,073,839 | 15,405,970 | 13.34 | 12,329,458 | 12,266,262 | 10.79 |
| 15,609,648 | 14,894,910 | 12.94 | 11,674,119 | 11,608,965 | 10.25 |
| 12,247,155 | 11,729,231 | 10.11 | 9,031,087 | 8,986,659 | 7.89 |
| 10,907,012 | 10,448,871 | 9.05 | 8,141,347 | 8,100,188 | 7.14 |
| 11,973,966 | 11,482,501 | 9.95 | 9,093,261 | 9,048,379 | 7.96 |
| 17,252,361 | 16,463,403 | 14.32 | 12,938,383 | 12,868,991 | 11.37 |
| 10,806,857 | 10,324,558 | 8.99 | 8,141,634 | 8,103,577 | 7.15 |
| 10,913,659 | 10,463,709 | 9.12 | 8,117,116 | 8,077,124 | 7.14 |
| 11,069,282 | 10,581,127 | 9.19 | 8,370,271 | 8,330,026 | 7.34 |
| 1,057,722 | 1,016,880 | 0.89 | 796,404 | 792,759 | 0.70 |
| 10,960,006 | 10,436,905 | 9.07 | 8,327,421 | 8,286,340 | 7.30 |
| 10,981,086 | 10,489,461 | 9.13 | 8,294,628 | 8,255,154 | 7.29 |
| 12,021,087 | 11,544,331 | 10.06 | 9,258,694 | 9,213,373 | 8.14 |
| 11,070,232 | 10,594,834 | 9.18 | 8,350,263 | 8,308,174 | 7.32 |
| 12,418,797 | 11,887,244 | 10.32 | 9,427,808 | 9,380,081 | 8.28 |
| 13,371,881 | 12,182,829 | 10.62 | 9,423,498 | 9,375,222 | 8.27 |
| 12,719,940 | 12,110,720 | 10.53 | 9,494,796 | 9,446,322 | 8.34 |
| 12,619,027 | 12,060,310 | 10.52 | 9,474,340 | 9,421,495 | 8.34 |
| 16,028,913 | 15,163,897 | 13.16 | 11,704,238 | 11,643,016 | 10.27 |
| 13,500,176 | 12,809,054 | 11.14 | 10,101,092 | 10,048,328 | 8.87 |
| 14,291,205 | 13,623,022 | 11.82 | 10,663,374 | 10,606,599 | 9.36 |
| 17,755,447 | 16,619,115 | 14.47 | 13,373,131 | 13,296,167 | 11.76 |
| 12,005,315 | 11,484,854 | 9.98 | 9,165,177 | 9,119,988 | 8.04 |
| 11,716,639 | 11,253,652 | 9.74 | 8,835,546 | 8,791,169 | 7.74 |
| 15,333,333 | 14,649,239 | 12.76 | 11,602,942 | 11,539,121 | 10.21 |
| 13,313,445 | 12,623,097 | 11.00 | 9,711,915 | 9,656,200 | 8.52 |
| 15,087,784 | 14,297,701 | 12.43 | 10,882,153 | 10,816,183 | 9.54 |
| 12,804,168 | 12,165,298 | 10.57 | 9,463,249 | 9,413,764 | 8.31 |
| 12,198,603 | 11,706,831 | 10.14 | 9,276,316 | 9,232,191 | 8.12 |
| 12,605,109 | 12,052,758 | 10.48 | 9,607,231 | 9,556,346 | 8.44 |
| 10,703,491 | 10,252,231 | 8.90 | 7,996,967 | 7,958,579 | 7.00 |
| 6,078,228 | 5,839,512 | 5.07 | 4,615,696 | 4,594,964 | 4.04 |
| 10,759,204 | 10,287,491 | 8.93 | 8,163,087 | 8,122,809 | 7.16 |
| 11,061,631 | 10,607,875 | 9.21 | 8,188,561 | 8,145,636 | 7.20 |
| 12,401,619 | 11,800,748 | 10.26 | 9,371,629 | 9,325,778 | 8.22 |
| 11,921,755 | 11,393,328 | 9.87 | 8,998,119 | 8,951,803 | 7.89 |
| 12,924,225 | 12,350,467 | 10.77 | 9,733,120 | 9,684,583 | 8.55 |
| 11,775,684 | 11,301,921 | 9.82 | 9,018,313 | 8,975,114 | 7.91 |
| 10,749,449 | 10,147,512 | 8.84 | 7,319,953 | 7,271,745 | 6.43 |
| 12,436,081 | 11,893,275 | 10.36 | 9,198,505 | 9,149,475 | 8.08 |
| 13,731,246 | 13,134,502 | 11.43 | 10,199,151 | 10,147,924 | 8.95 |
| 13,069,182 | 12,535,235 | 10.89 | 9,923,707 | 9,872,736 | 8.71 |
| 18,600,212 | 17,760,696 | 15.38 | 13,836,700 | 13,765,302 | 12.13 |
| 11,088,508 | 10,576,009 | 9.18 | 8,014,385 | 7,972,980 | 7.03 |
| 12,378,789 | 11,867,465 | 10.31 | 9,099,520 | 9,053,167 | 7.99 |
| 15,067,549 | 14,404,874 | 12.52 | 11,304,402 | 11,242,903 | 9.92 |
| 12,515,418 | 11,980,431 | 10.42 | 9,569,913 | 9,520,858 | 8.40 |
| 143,986 | 139,136 | 0.12 | 106,865 | 106,340 | 0.09 |
| 17,412,721 | 16,501,457 | 14.26 | 12,917,937 | 12,843,221 | 11.30 |
| 12,997,996 | 12,331,202 | 10.68 | 9,640,572 | 9,588,392 | 8.46 |
| 14,996,072 | 14,279,609 | 12.36 | 11,094,544 | 11,032,388 | 9.71 |
| 11,026,120 | 10,430,449 | 9.08 | 7,536,229 | 7,485,955 | 6.61 |
| 12,536,912 | 12,023,216 | 10.44 | 9,441,331 | 9,393,191 | 8.30 |
| 6,863,454 | 6,432,839 | 5.59 | 4,384,393 | 4,358,401 | 3.84 |
| 10,848,382 | 10,398,937 | 9.01 | 8,094,965 | 8,054,858 | 7.09 |
| 13,518,050 | 12,956,962 | 11.24 | 10,244,752 | 10,193,611 | 8.98 |
| 12,435,544 | 11,828,492 | 10.18 | 9,235,973 | 9,189,532 | 8.03 |
| 11,529,839 | 11,094,695 | 9.57 | 8,944,844 | 8,901,671 | 7.82 |
| 13,201,448 | 12,585,846 | 10.95 | 10,077,513 | 10,023,660 | 8.86 |
| 11,839,473 | 11,376,272 | 9.86 | 8,908,763 | 8,860,102 | 7.82 |
| 6,860,487 | 6,459,367 | 5.61 | 4,478,265 | 4,451,818 | 3.92 |
| 10,363,606 | 9,809,624 | 8.56 | 7,021,531 | 6,977,921 | 6.17 |
| 10,657,720 | 10,236,527 | 8.90 | 8,051,251 | 8,011,587 | 7.07 |
| 9,198,269 | 8,795,421 | 7.67 | 6,691,839 | 6,655,571 | 5.89 |
| 12,225,244 | 11,639,493 | 10.09 | 9,023,817 | 8,974,299 | 7.92 |
| 11,662,428 | 11,102,277 | 9.62 | 8,693,353 | 8,648,384 | 7.61 |
| 12,989,393 | 12,206,757 | 10.63 | 9,662,274 | 9,612,292 | 8.50 |
| 12,176,438 | 11,664,612 | 10.13 | 9,502,782 | 9,456,105 | 8.34 |
| 11,715,316 | 11,216,265 | 9.73 | 8,658,038 | 8,611,329 | 7.59 |
| 14,533,788 | 13,752,973 | 11.96 | 10,602,094 | 10,538,536 | 9.31 |
| 12,138,605 | 11,566,883 | 10.06 | 8,781,736 | 8,731,146 | 7.72 |
| 12,035,817 | 11,481,361 | 9.96 | 8,788,926 | 8,740,613 | 7.70 |
| 17,710,426 | 16,914,030 | 14.73 | 13,996,814 | 13,935,690 | 12.32 |
| 11,778,630 | 11,279,862 | 9.77 | 8,819,860 | 8,774,196 | 7.74 |
| 11,987,904 | 11,427,213 | 9.87 | 8,943,690 | 8,899,459 | 7.82 |
| 10,448,610 | 9,977,120 | 8.65 | 7,898,648 | 7,861,215 | 6.92 |
| 10,431,838 | 9,906,286 | 8.56 | 7,764,456 | 7,728,708 | 6.79 |
| 13,256,545 | 12,562,557 | 10.97 | 9,851,467 | 9,798,684 | 8.67 |
| 13,983,395 | 13,287,316 | 11.57 | 10,308,770 | 10,259,002 | 9.06 |
| 12,985,922 | 12,396,095 | 10.77 | 9,592,485 | 9,540,314 | 8.43 |
| 12,326,905 | 11,773,978 | 10.19 | 9,248,871 | 9,203,403 | 8.10 |
| 12,446,359 | 11,879,732 | 10.34 | 9,233,086 | 9,180,766 | 8.12 |
| 11,647,756 | 11,040,817 | 9.55 | 8,523,194 | 8,477,193 | 7.46 |
| 13,178,610 | 11,896,797 | 10.36 | 9,271,263 | 9,225,165 | 8.15 |
| 11,483,934 | 10,932,251 | 9.53 | 8,969,076 | 8,926,151 | 7.87 |
| 13,938,041 | 13,336,158 | 11.62 | 10,656,851 | 10,603,350 | 9.36 |
| 11,526,261 | 10,998,601 | 9.52 | 8,682,078 | 8,638,080 | 7.60 |
| 15,745,062 | 15,053,383 | 13.08 | 12,173,699 | 12,112,129 | 10.67 |
| 9,965,923 | 9,545,106 | 8.25 | 7,478,586 | 7,442,718 | 6.54 |
| 11,653,929 | 11,168,018 | 9.72 | 9,060,087 | 9,018,093 | 7.95 |
| 11,635,455 | 11,058,525 | 9.60 | 8,342,893 | 8,300,311 | 7.33 |
| 11,646,297 | 11,148,438 | 9.66 | 8,400,207 | 8,358,608 | 7.37 |
| 10,748,614 | 10,327,897 | 8.99 | 8,036,443 | 7,997,274 | 7.07 |
| 10,568,382 | 10,105,762 | 8.75 | 7,889,680 | 7,850,830 | 6.91 |
| 11,841,622 | 11,382,107 | 9.85 | 8,943,548 | 8,898,921 | 7.83 |
| 11,691,136 | 11,146,667 | 9.63 | 9,041,574 | 9,001,370 | 7.88 |
| 14,493,451 | 13,823,722 | 12.04 | 10,868,519 | 10,807,667 | 9.56 |
| 12,616,232 | 12,069,098 | 10.47 | 9,599,745 | 9,547,188 | 8.41 |
| 11,732,242 | 11,062,372 | 9.65 | 8,187,354 | 8,138,527 | 7.19 |
| 10,275,603 | 9,853,420 | 8.52 | 7,725,688 | 7,687,177 | 6.76 |
| 10,953,576 | 10,518,955 | 9.12 | 8,255,795 | 8,213,949 | 7.23 |
| 3,995,031 | 3,697,482 | 3.22 | 2,412,314 | 2,395,717 | 2.12 |
| 11,211,632 | 10,691,895 | 9.29 | 8,208,648 | 8,169,133 | 7.21 |
| 13,572,589 | 12,967,546 | 11.23 | 10,225,148 | 10,173,895 | 8.94 |
| 6,627,616 | 6,360,941 | 5.50 | 5,008,835 | 4,984,721 | 4.38 |
| 11,764,013 | 11,258,119 | 9.76 | 8,827,600 | 8,782,088 | 7.75 |
| 2,401,470 | 2,315,754 | 2.00 | 1,769,941 | 1,762,209 | 1.54 |
| 9,797,113 | 9,352,026 | 8.10 | 7,228,974 | 7,192,540 | 6.34 |
| 10,422,469 | 9,931,609 | 8.66 | 7,552,629 | 7,508,231 | 6.63 |
| 13,908,357 | 13,335,312 | 11.63 | 10,753,356 | 10,700,615 | 9.45 |
| 15,789,591 | 15,073,326 | 13.12 | 12,161,493 | 12,097,619 | 10.68 |
| 8,840,784 | 8,332,742 | 7.26 | 6,259,122 | 6,225,006 | 5.49 |
| 10,674,812 | 10,236,848 | 8.87 | 8,156,722 | 8,117,549 | 7.16 |
| 12,436,324 | 11,840,569 | 10.33 | 9,496,676 | 9,450,143 | 8.36 |
| 13,006,242 | 12,338,133 | 10.73 | 9,418,446 | 9,368,446 | 8.28 |
| 10,865,991 | 10,360,084 | 8.96 | 7,903,591 | 7,866,419 | 6.92 |
| 13,261,267 | 12,720,129 | 11.08 | 10,012,708 | 9,962,362 | 8.81 |
| 11,664,035 | 11,209,448 | 9.68 | 9,063,344 | 9,020,820 | 7.92 |
| 12,683,898 | 12,176,713 | 10.47 | 9,731,101 | 9,683,103 | 8.48 |
| 10,678,514 | 10,207,393 | 8.81 | 7,946,350 | 7,905,669 | 6.95 |
| 7,708,253 | 7,249,250 | 6.33 | 5,319,777 | 5,282,599 | 4.67 |
| 14,849,629 | 14,175,720 | 12.33 | 11,110,038 | 11,048,787 | 9.76 |
| 12,027,830 | 11,362,881 | 9.93 | 8,535,693 | 8,478,971 | 7.51 |
| 5,974,983 | 5,466,743 | 4.76 | 3,928,522 | 3,901,771 | 3.45 |
| 10,650,776 | 10,024,119 | 8.71 | 7,372,998 | 7,328,256 | 6.46 |
| 7,740,489 | 7,291,414 | 6.35 | 5,198,751 | 5,165,999 | 4.56 |
| 13,370,489 | 12,814,664 | 11.18 | 10,125,245 | 10,075,324 | 8.92 |
| 9,583,975 | 9,125,635 | 7.95 | 7,005,108 | 6,969,916 | 6.15 |
| 13,197,317 | 12,613,463 | 10.99 | 10,148,285 | 10,098,677 | 8.92 |
| 12,568,291 | 12,018,260 | 10.44 | 9,529,089 | 9,480,901 | 8.37 |
| 11,249,891 | 10,782,325 | 9.37 | 8,614,940 | 8,568,829 | 7.58 |
| 1,736,551 | 1,668,610 | 1.45 | 1,305,802 | 1,299,531 | 1.15 |
| 9,231,628 | 8,427,334 | 7.33 | 6,238,597 | 6,199,001 | 5.47 |
| 8,926,888 | 8,336,881 | 7.28 | 6,165,457 | 6,128,511 | 5.42 |
| 10,345,263 | 9,458,038 | 8.26 | 6,990,260 | 6,943,576 | 6.14 |
| 8,821,857 | 8,064,309 | 7.03 | 6,082,909 | 6,048,965 | 5.34 |
| 12,339,102 | 11,612,150 | 10.13 | 9,508,367 | 9,463,085 | 8.36 |
| 12,285,187 | 11,718,564 | 10.18 | 9,318,477 | 9,271,055 | 8.20 |
| 12,819,912 | 11,807,600 | 10.24 | 9,229,736 | 9,184,223 | 8.08 |
| 11,740,190 | 11,196,315 | 9.72 | 8,565,136 | 8,520,003 | 7.52 |
| 11,614,810 | 11,001,944 | 9.61 | 8,434,074 | 8,389,894 | 7.43 |
| 12,668,585 | 12,146,536 | 10.58 | 9,606,041 | 9,559,066 | 8.44 |
| 12,678,357 | 12,164,523 | 10.59 | 9,716,856 | 9,670,572 | 8.54 |
| 12,328,811 | 11,774,281 | 10.22 | 9,226,320 | 9,176,416 | 8.09 |
| 7,041,324 | 6,513,036 | 5.67 | 4,697,232 | 4,666,611 | 4.12 |
| 15,045,607 | 14,395,726 | 12.52 | 11,348,006 | 11,286,125 | 9.97 |
| 13,195,352 | 12,635,660 | 10.98 | 9,929,082 | 9,875,805 | 8.72 |
| 18,166,834 | 17,357,578 | 15.07 | 13,478,294 | 13,399,500 | 11.83 |
| 8,424,272 | 7,870,832 | 6.84 | 5,557,744 | 5,521,432 | 4.87 |
| 11,463,552 | 10,912,475 | 9.48 | 8,430,562 | 8,387,832 | 7.40 |
| 11,415,722 | 10,968,617 | 9.48 | 8,472,252 | 8,432,846 | 7.41 |
| 10,721,502 | 10,261,297 | 8.86 | 7,781,361 | 7,743,483 | 6.80 |
| 15,107,716 | 14,450,804 | 12.58 | 11,286,415 | 11,228,296 | 9.92 |
| 13,663,370 | 13,064,589 | 11.37 | 10,360,553 | 10,308,389 | 9.11 |
| 11,781,373 | 11,303,029 | 9.80 | 9,083,395 | 9,038,823 | 7.96 |
| 12,501,391 | 11,991,405 | 10.45 | 9,567,471 | 9,518,160 | 8.43 |
| 9,770,541 | 9,207,740 | 8.02 | 6,746,995 | 6,705,439 | 5.92 |
| 11,988,940 | 11,372,905 | 9.85 | 8,996,775 | 8,949,682 | 7.89 |
| 9,731,719 | 9,237,748 | 8.03 | 6,807,607 | 6,767,841 | 5.97 |
| 10,395,554 | 9,903,040 | 8.64 | 7,551,366 | 7,507,251 | 6.64 |
| 11,302,886 | 10,785,665 | 9.38 | 8,442,426 | 8,396,587 | 7.42 |
| 11,615,392 | 10,720,150 | 9.36 | 8,458,386 | 8,413,783 | 7.45 |
| 13,157,419 | 12,505,316 | 10.93 | 9,719,942 | 9,663,082 | 8.56 |
| 12,204,241 | 11,702,517 | 10.17 | 9,317,466 | 9,268,767 | 8.18 |
| 11,451,896 | 10,981,278 | 9.54 | 8,541,072 | 8,499,124 | 7.51 |
| 12,686,698 | 12,108,515 | 10.43 | 9,059,172 | 9,006,911 | 7.90 |
| 14,941,999 | 14,248,144 | 12.40 | 10,911,894 | 10,843,063 | 9.59 |
| 11,308,660 | 10,836,539 | 9.39 | 8,223,433 | 8,178,383 | 7.21 |
| 17,894,228 | 17,010,556 | 14.80 | 13,386,922 | 13,313,236 | 11.77 |
| 13,046,978 | 12,442,626 | 10.80 | 9,794,650 | 9,743,912 | 8.59 |
| 13,957,245 | 13,302,052 | 11.51 | 10,241,180 | 10,186,956 | 8.97 |
| 12,763,549 | 12,159,370 | 10.46 | 9,431,722 | 9,384,352 | 8.19 |
| 12,136,799 | 11,620,696 | 10.08 | 9,037,989 | 8,992,687 | 7.92 |
| 11,949,027 | 11,428,810 | 9.83 | 9,132,246 | 9,087,070 | 7.92 |
| 11,702,614 | 11,175,617 | 9.70 | 8,725,690 | 8,682,521 | 7.68 |
| 13,558,019 | 12,973,543 | 11.19 | 9,951,575 | 9,901,140 | 8.72 |
| 15,295,590 | 14,615,585 | 12.69 | 11,337,881 | 11,277,026 | 9.98 |
| 14,674,025 | 13,910,357 | 12.09 | 10,592,556 | 10,535,523 | 9.29 |
| 10,697,649 | 10,233,396 | 8.83 | 7,869,539 | 7,831,672 | 6.89 |
| 13,476,442 | 12,932,853 | 11.17 | 10,154,149 | 10,103,914 | 8.91 |
| 12,667,435 | 11,690,972 | 10.18 | 8,019,580 | 7,956,418 | 7.04 |
| 11,779,772 | 11,001,816 | 9.59 | 7,877,723 | 7,826,141 | 6.92 |
| 11,432,243 | 10,941,301 | 9.51 | 8,612,114 | 8,571,886 | 7.57 |
| 9,323,612 | 8,845,182 | 7.70 | 6,392,825 | 6,354,846 | 5.61 |
| 9,359,940 | 8,862,664 | 7.73 | 6,524,433 | 6,485,245 | 5.73 |
| 11,419,220 | 10,879,901 | 9.49 | 8,159,339 | 8,112,511 | 7.17 |

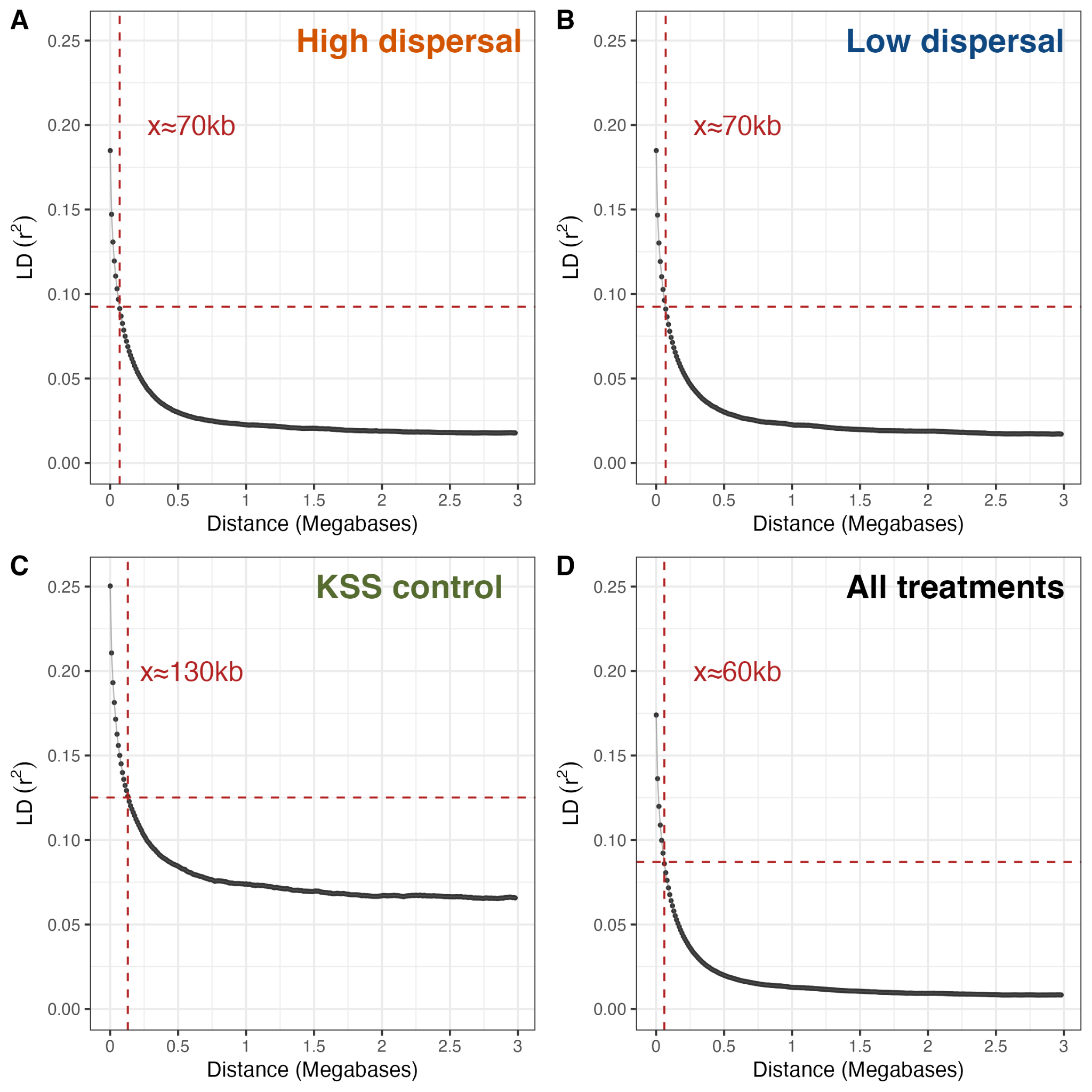

**Figure S1.** Linkage decay estimated from genetic samples derived from an experiment exposing *Tribolium castaneum* to artificial selection for dispersal propensity. Panels show each dispersal treatment and the ancestral line considered separately (A-C), and considering all samples as a single population (D). The value of x represented in red shows the genomic distance over which the linkage halves from the maximum estimated value, rounded to the nearest 10kb. Note that sample sizes differ across panels/treatments and that the pattern we observe is consistent with the estimates of linkage reducing with increasing sample size (A n = 93; B n = 96, C n = 18, D n = 207).

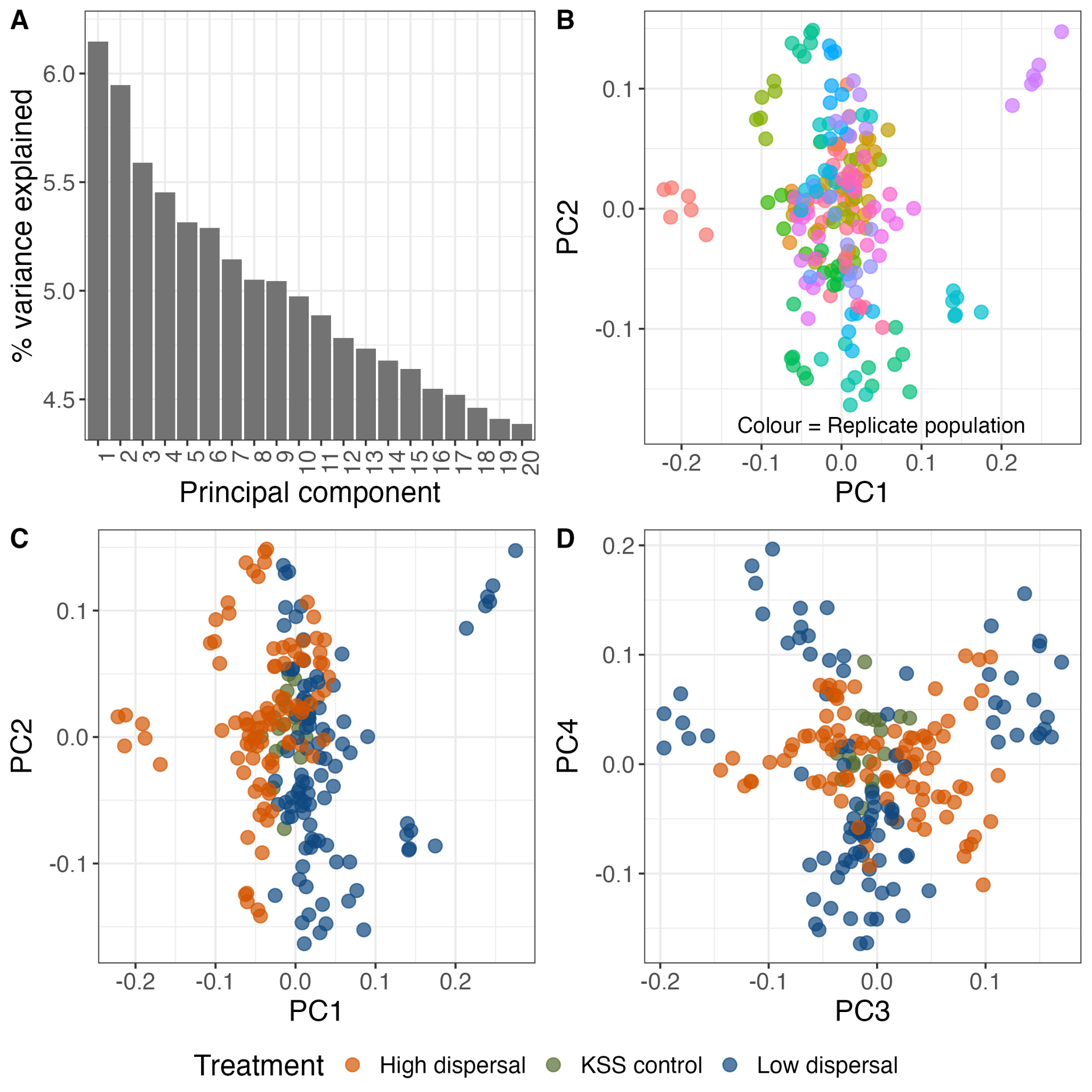

**Figure S2.** Principal component analysis (PCA) on genetic samples derived from an experiment exposing *Tribolium castaneum* to artificial selection for dispersal propensity. Panels show, **A**) the explanatory power of each PC, **B**) strong clustering of samples by their population of origin (colour), **C**) how PC1 largely represents dispersal selection treatment and **C&D**) how samples spread out from the core of the parameter space occupied by the ancestral KSS population. Interestingly, low dispersal samples appear to be less similar to KSS control samples than high dispersal samples are, in both directions, on both PC3 and PC4.

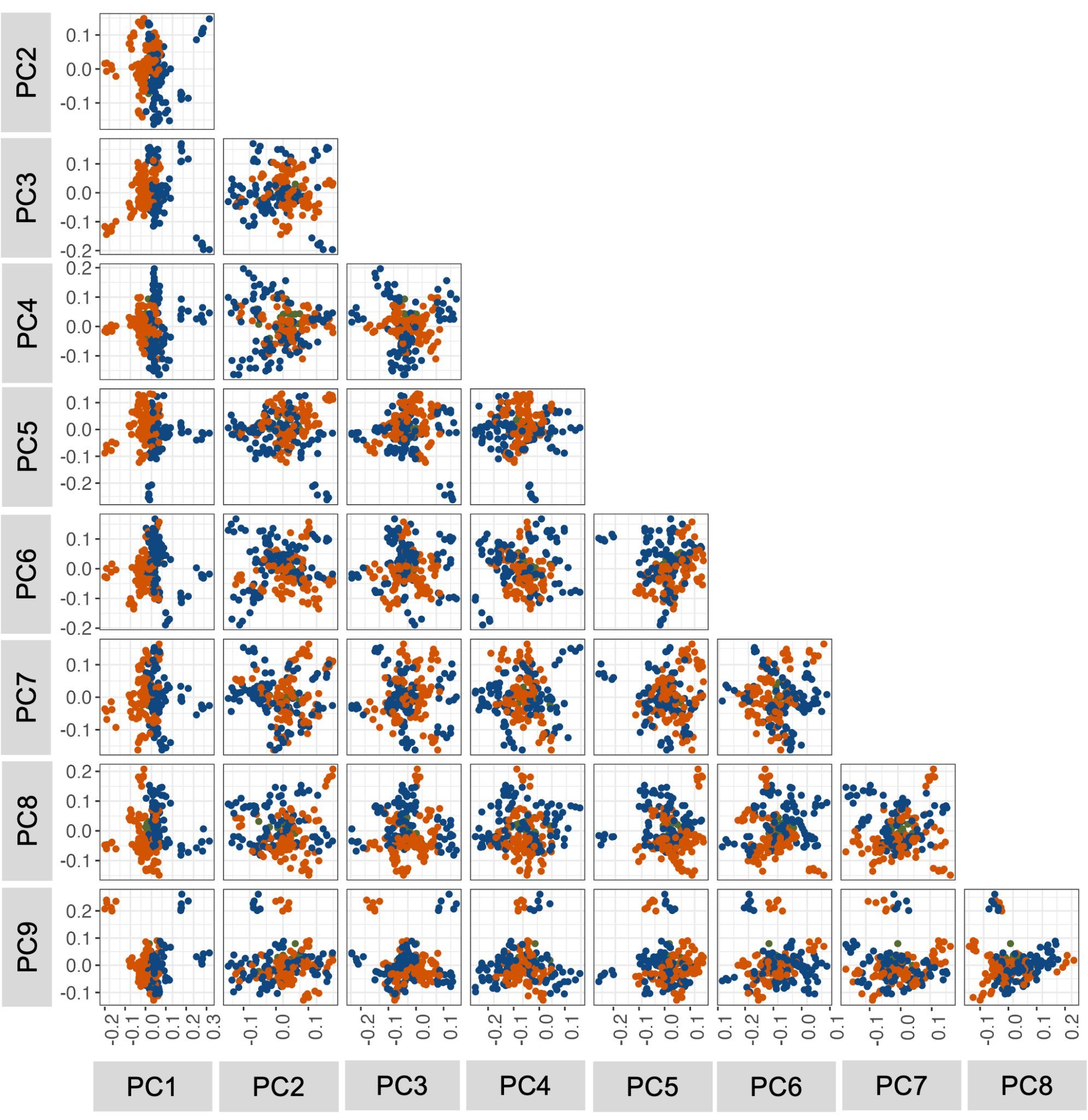

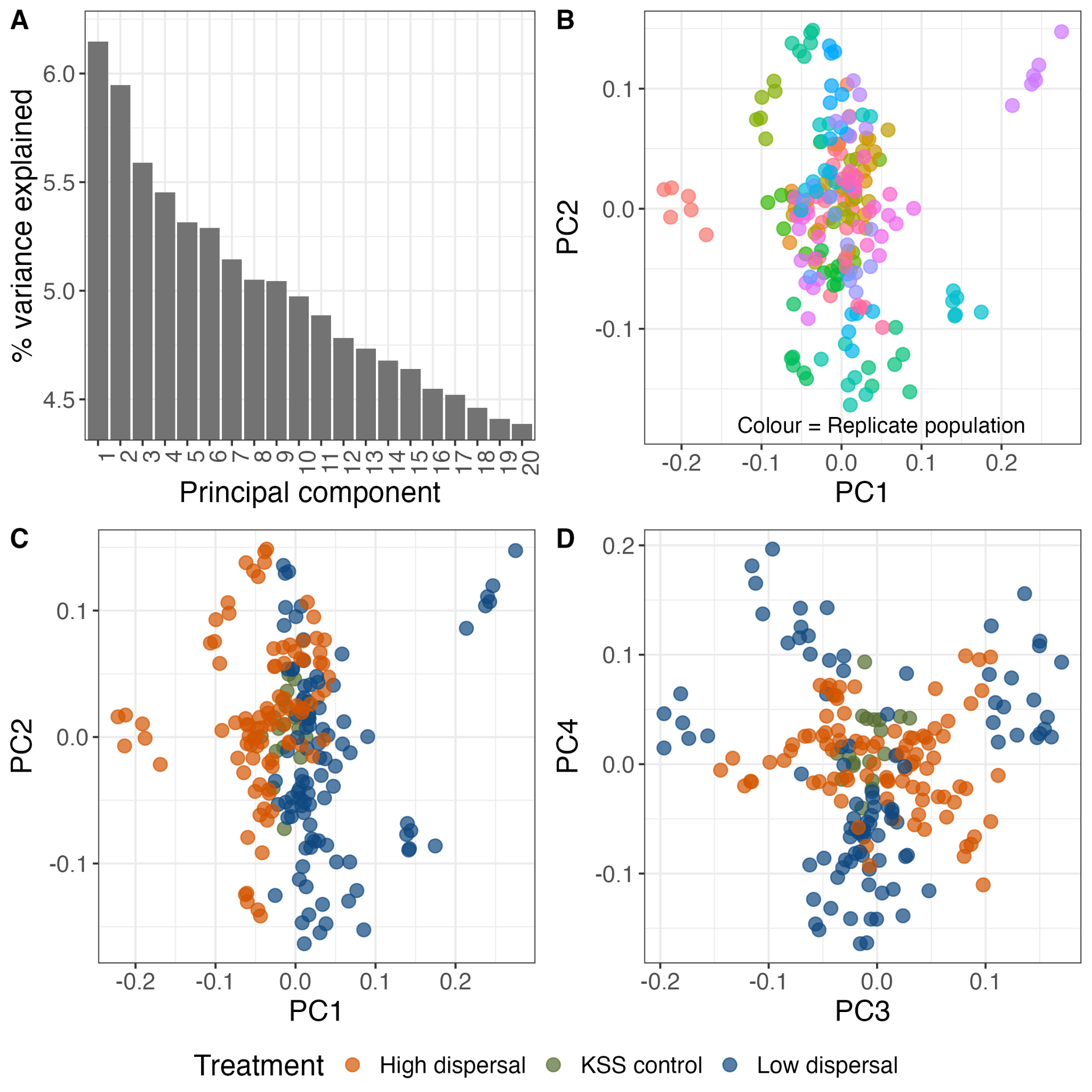

**Figure S3.** Matrix of PCs explaining >5% of total variance in a PCA analysis of genomic data derived from an experiment exposing *Tribolium castaneum* to artificial selection for dispersal propensity. PC1 appears to capture most of the difference between dispersal selection regimes.

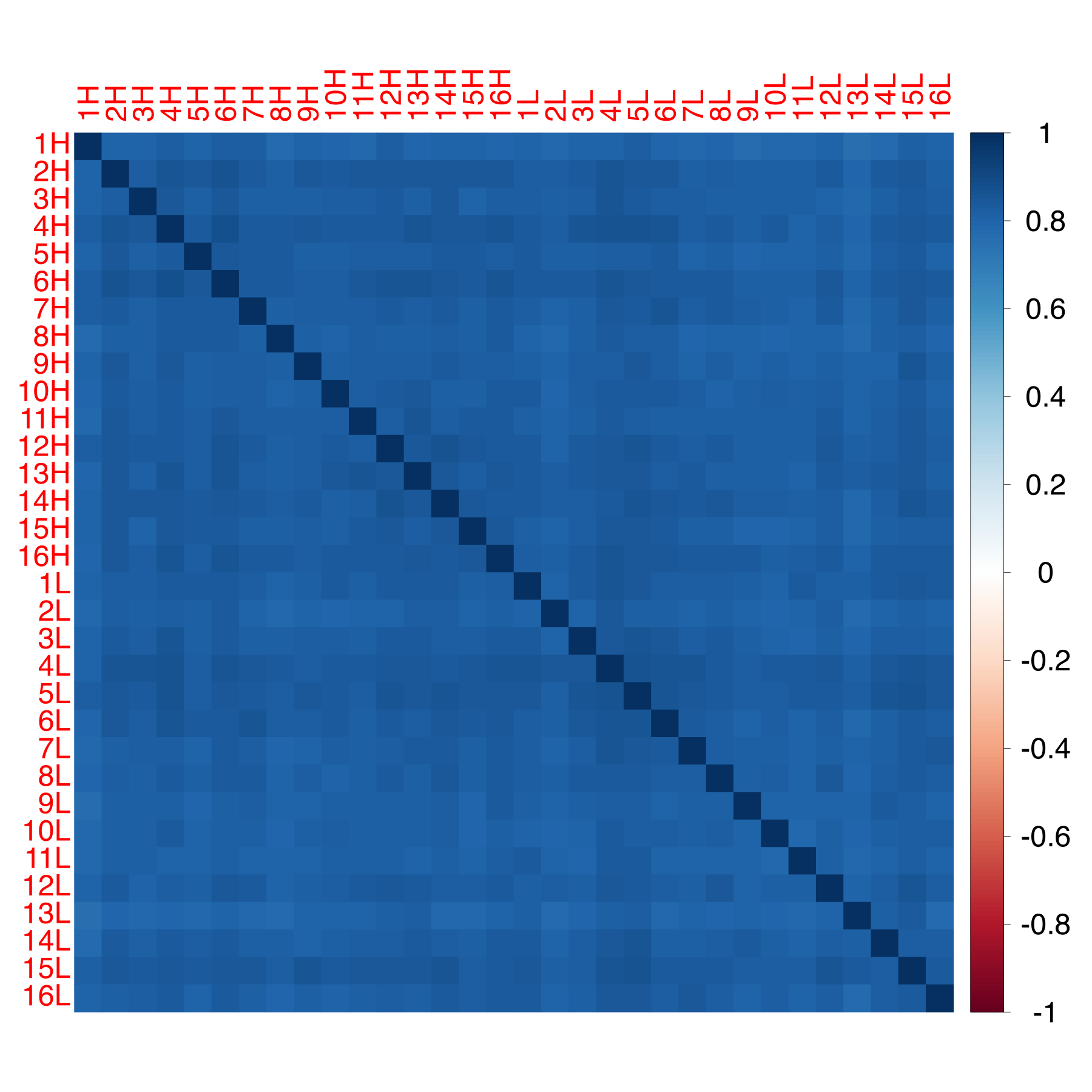

**Figure S4.** Visualisation of the background allele frequency matrix (omega) generated by BayPass, using a set of genome-wide SNPs derived from an experiment exposing *Tribolium castaneum* to artificial selection for dispersal propensity. Each row and column represents a single, independently evolving high dispersal (n=16) or low dispersal (n=16) population, with the colour of each square showing the correlation in allele frequencies between population pairs.

A)

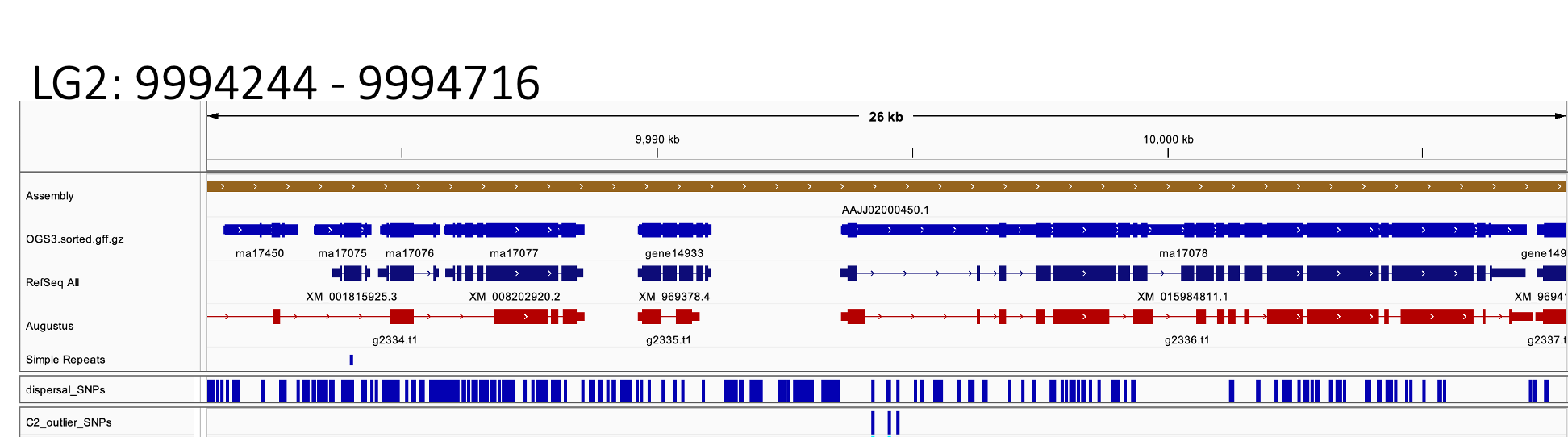

B)

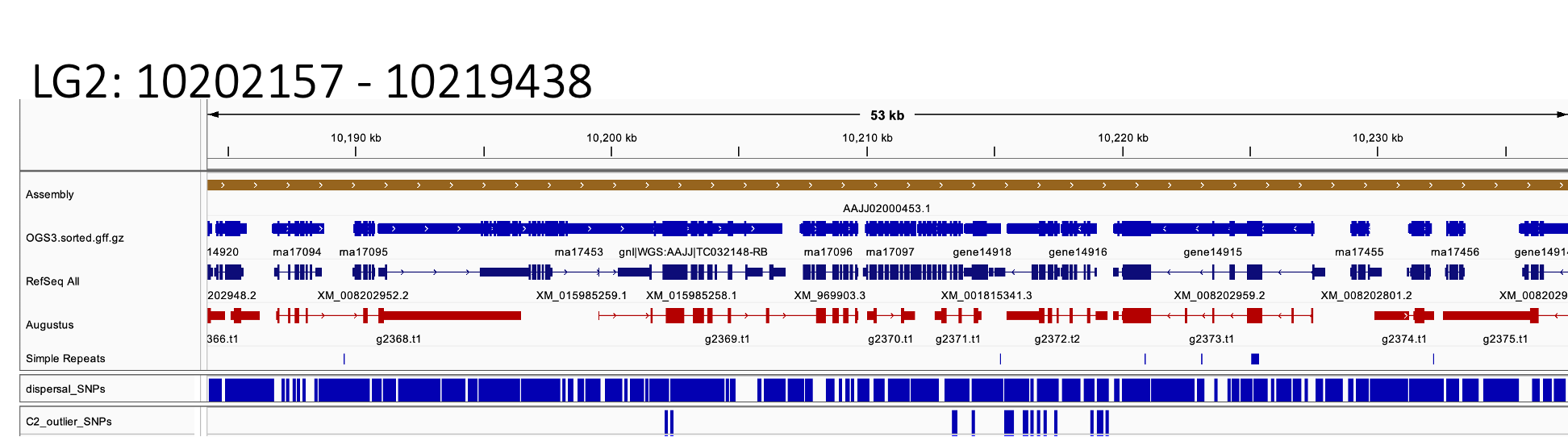

C)

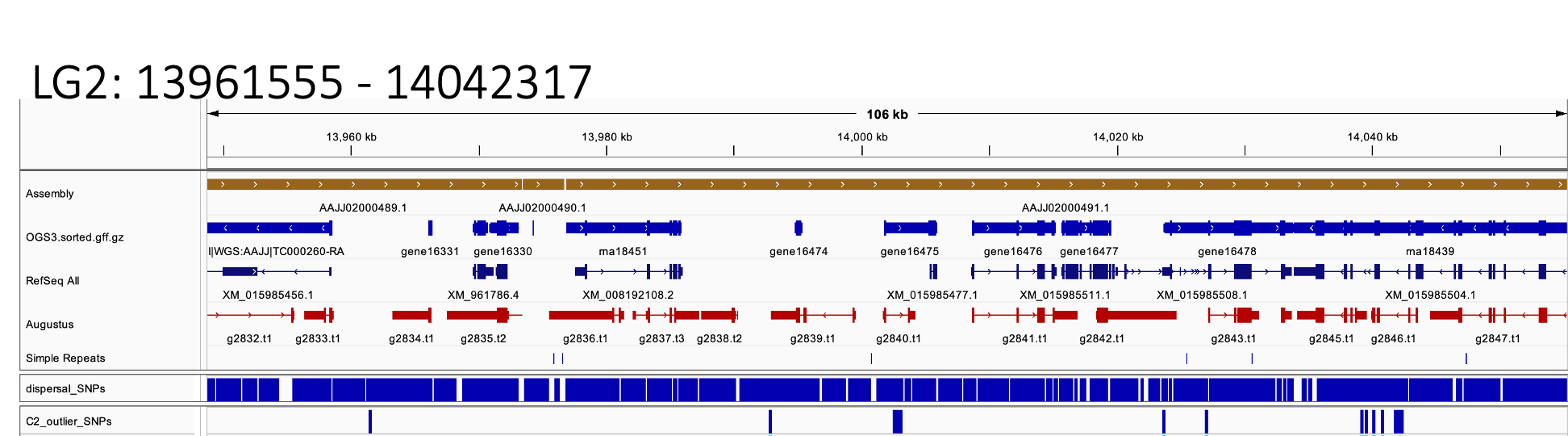

D)
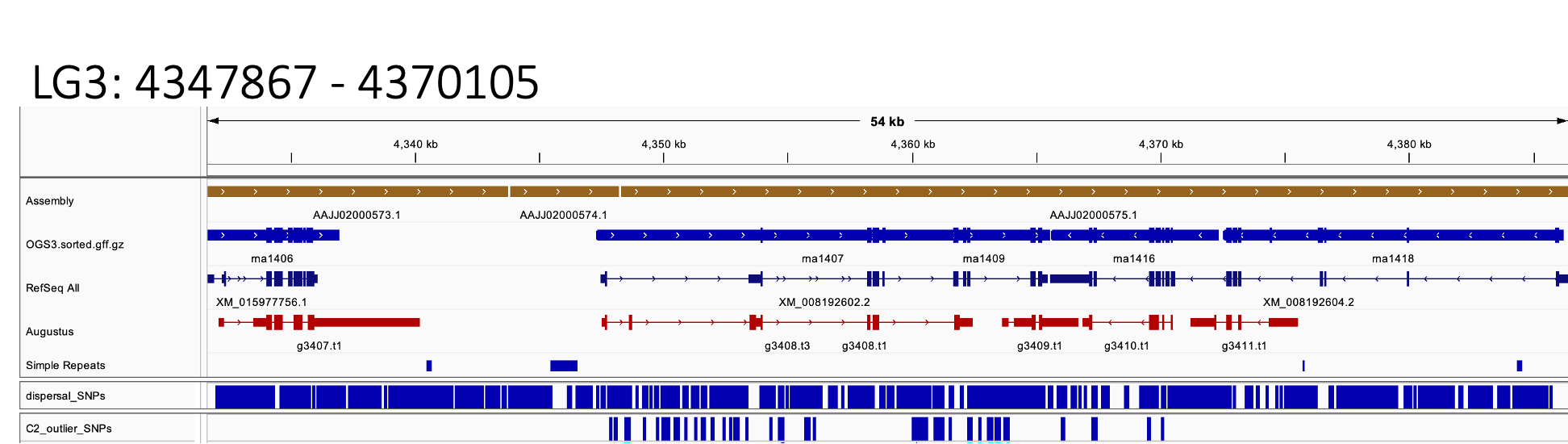

E)

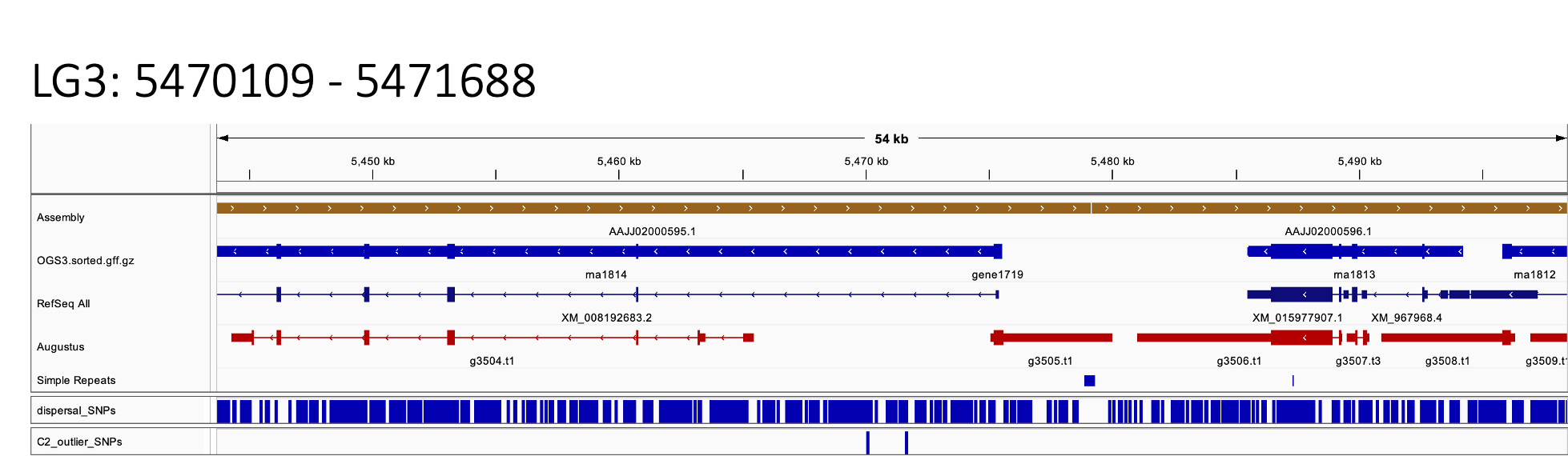

F)
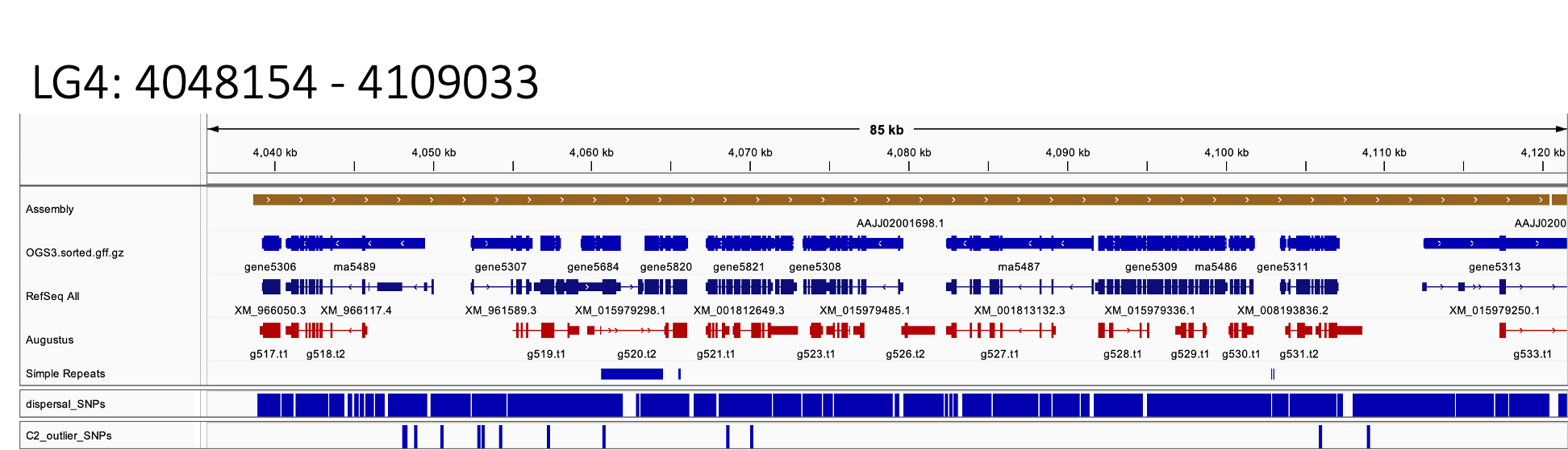

G)
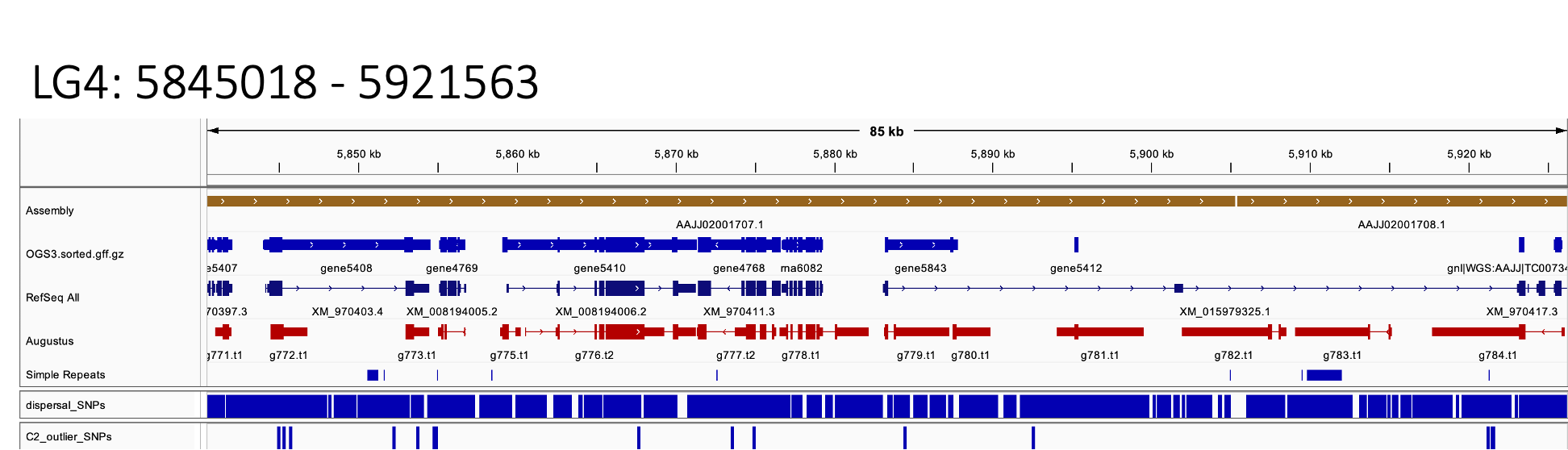

H)

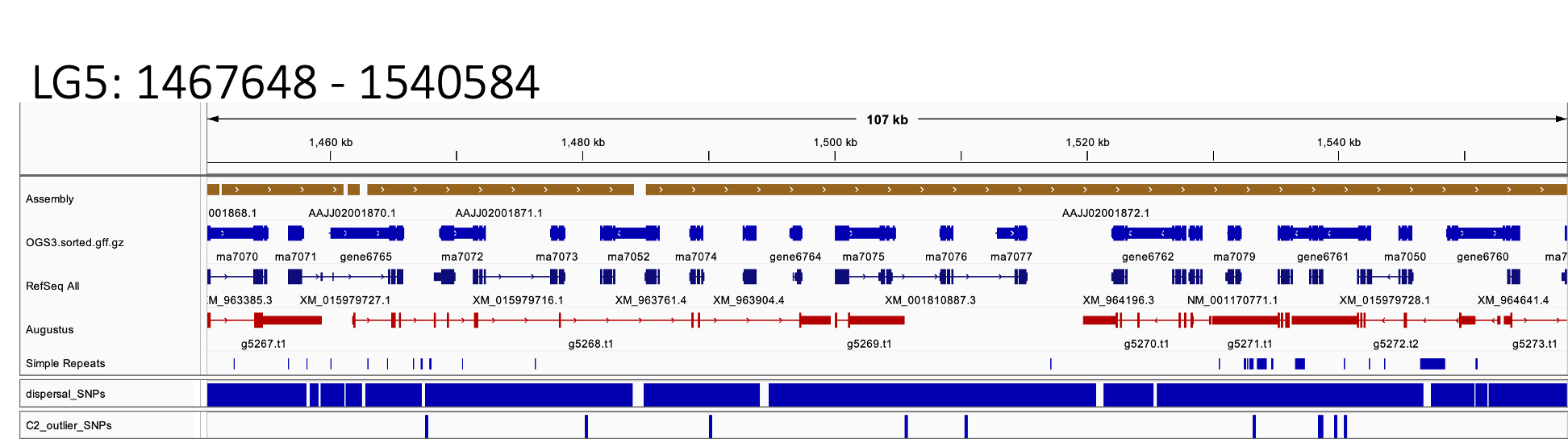

I)

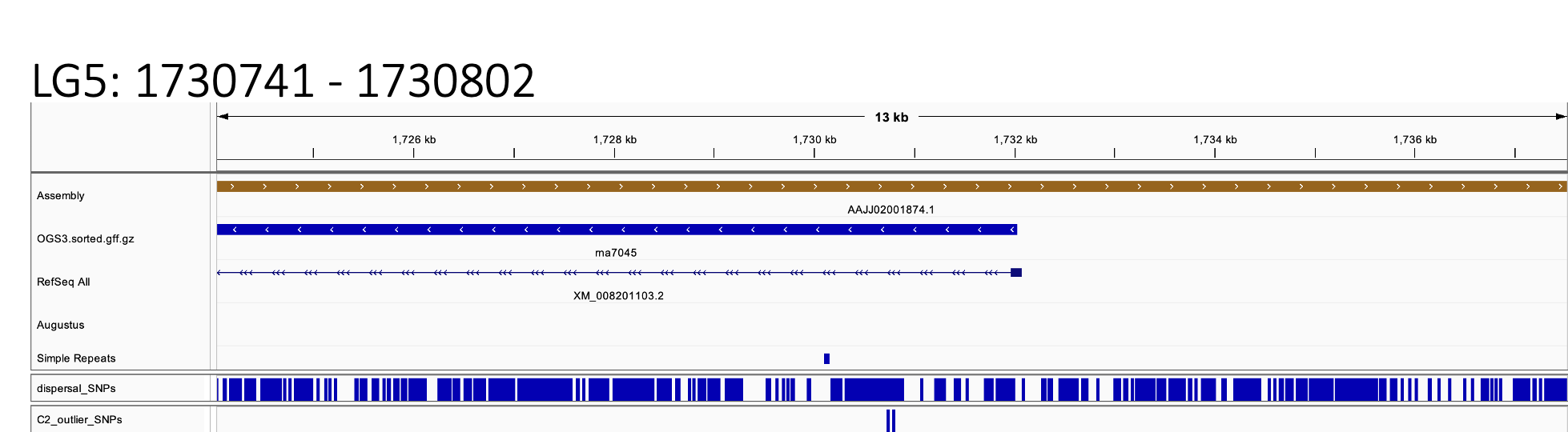

J)

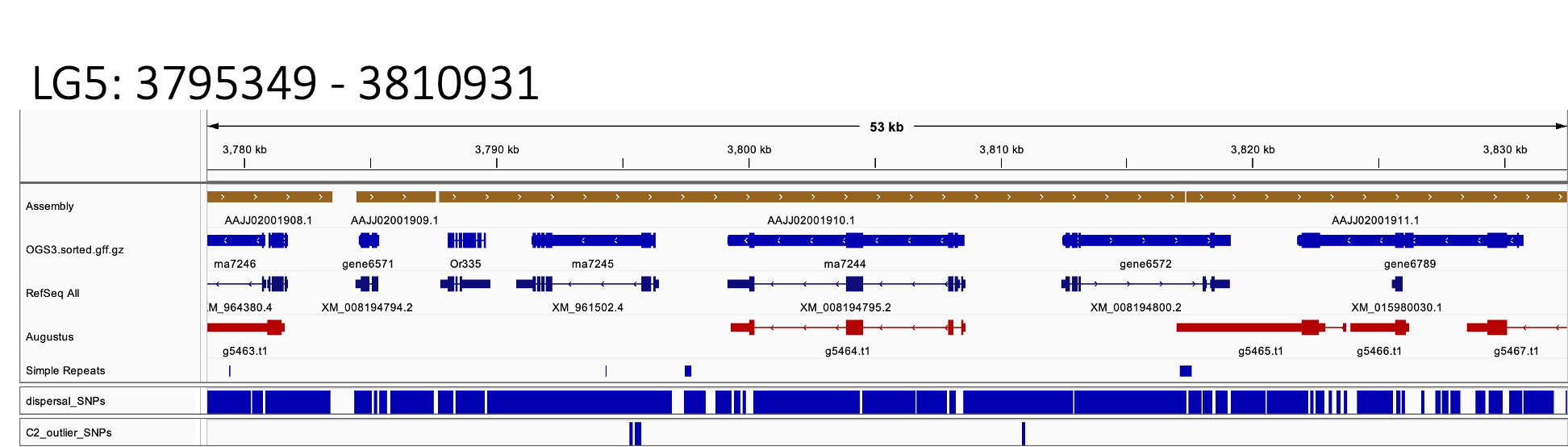

K)

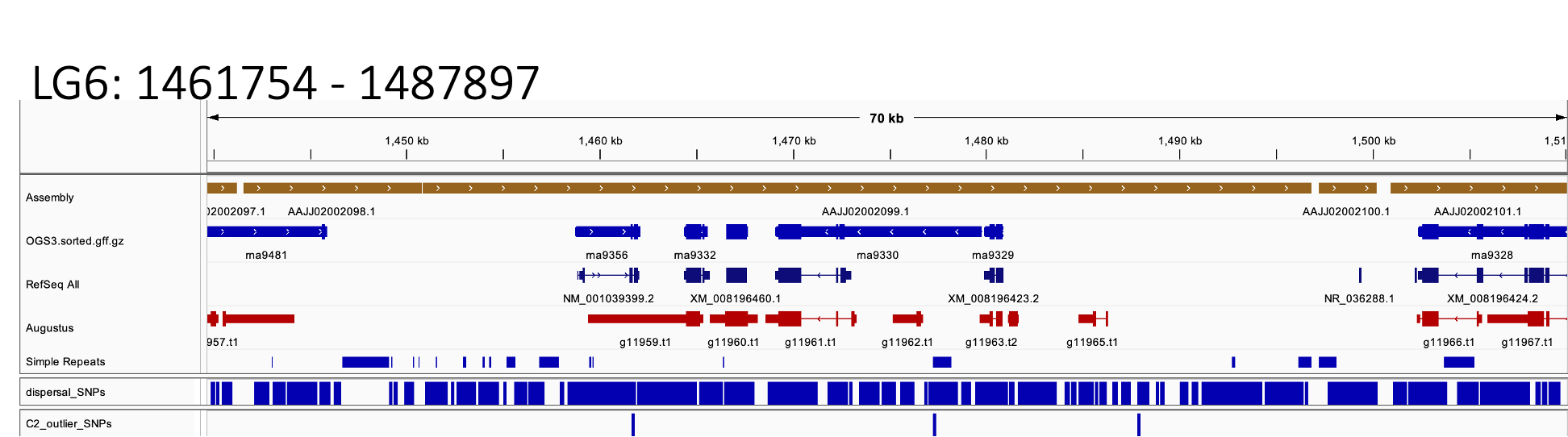

L)

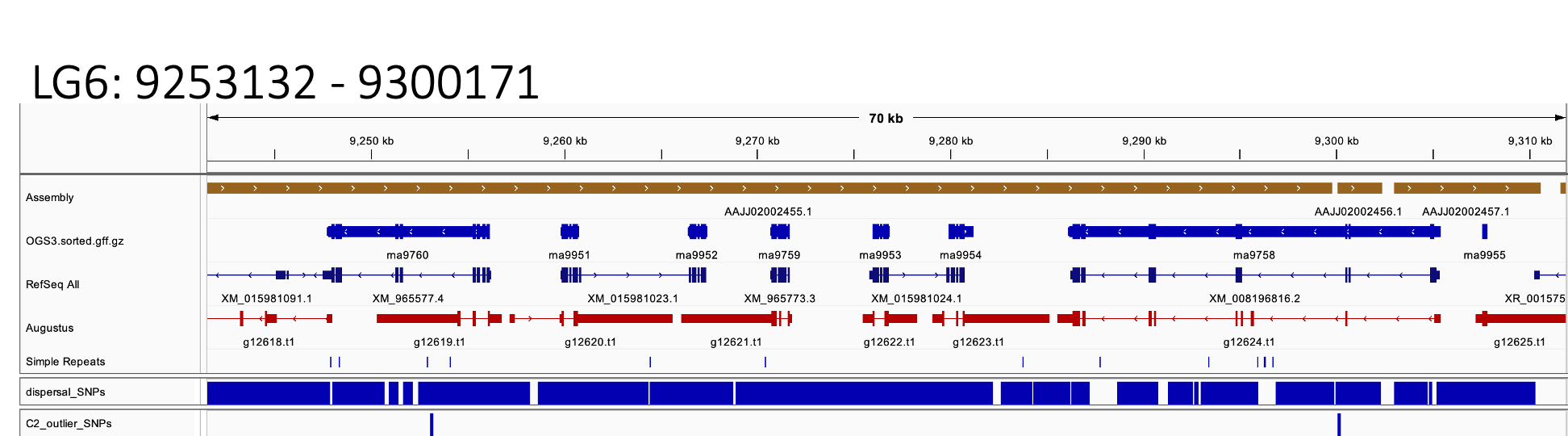

M)

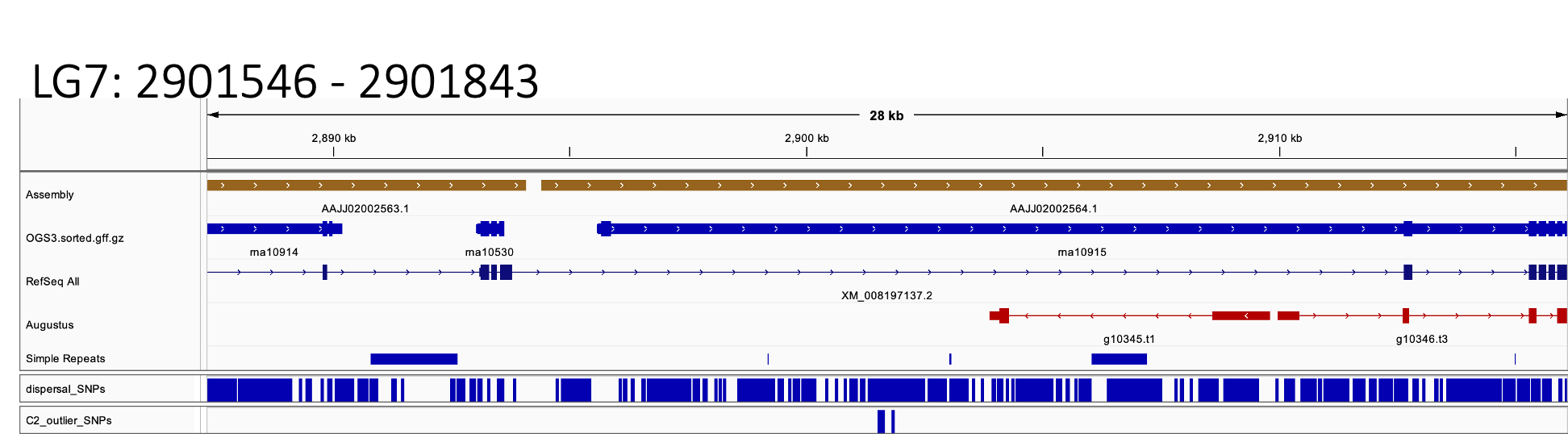

N)

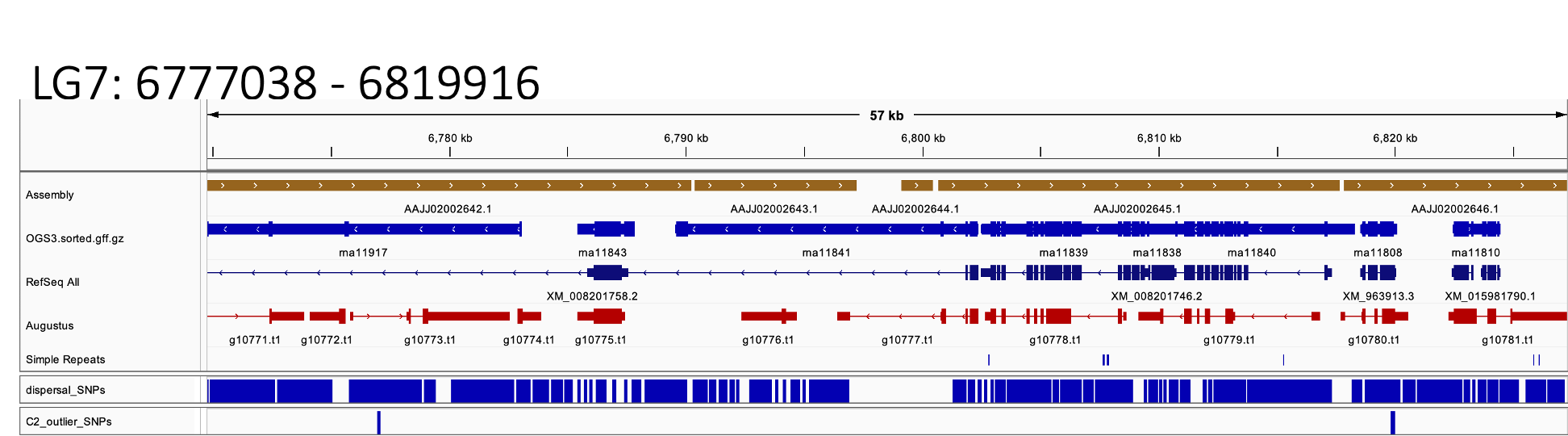

O)

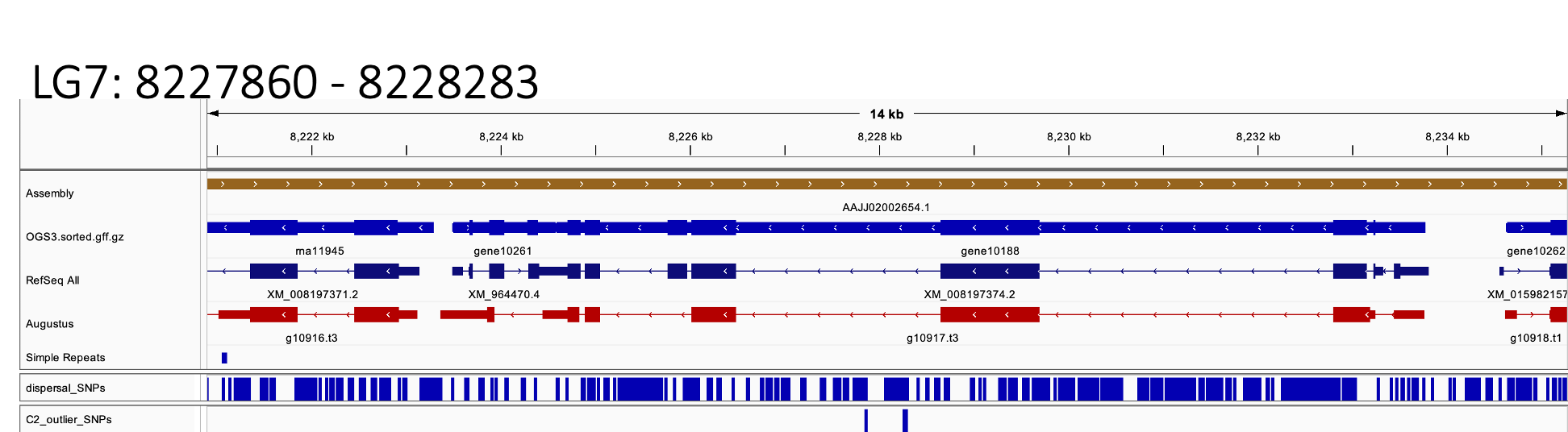

P)

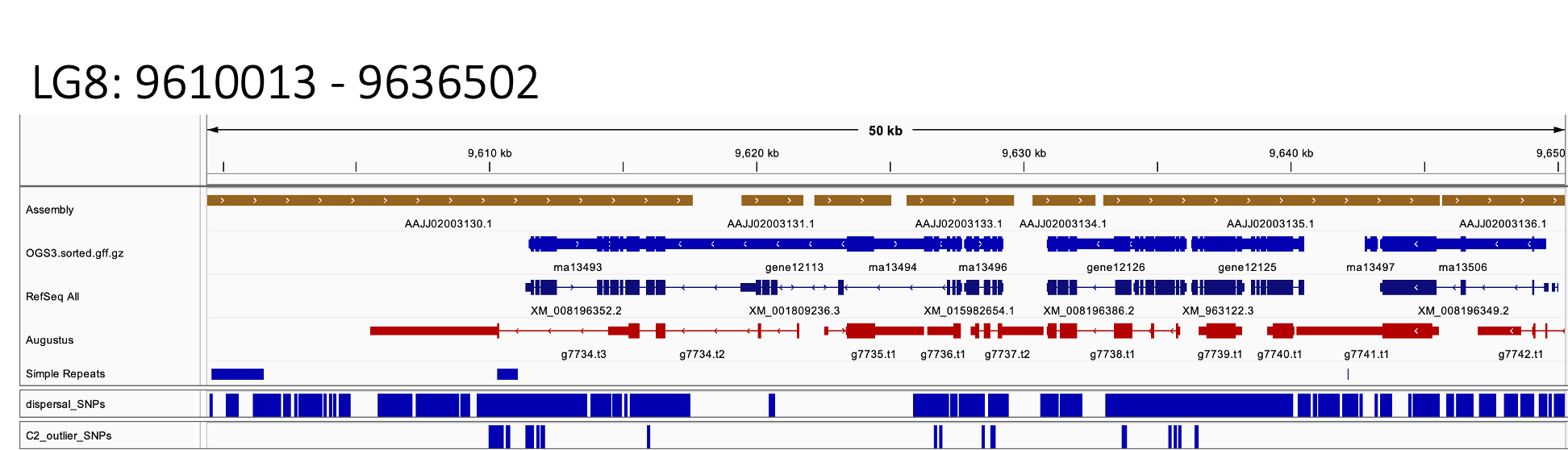

Q)

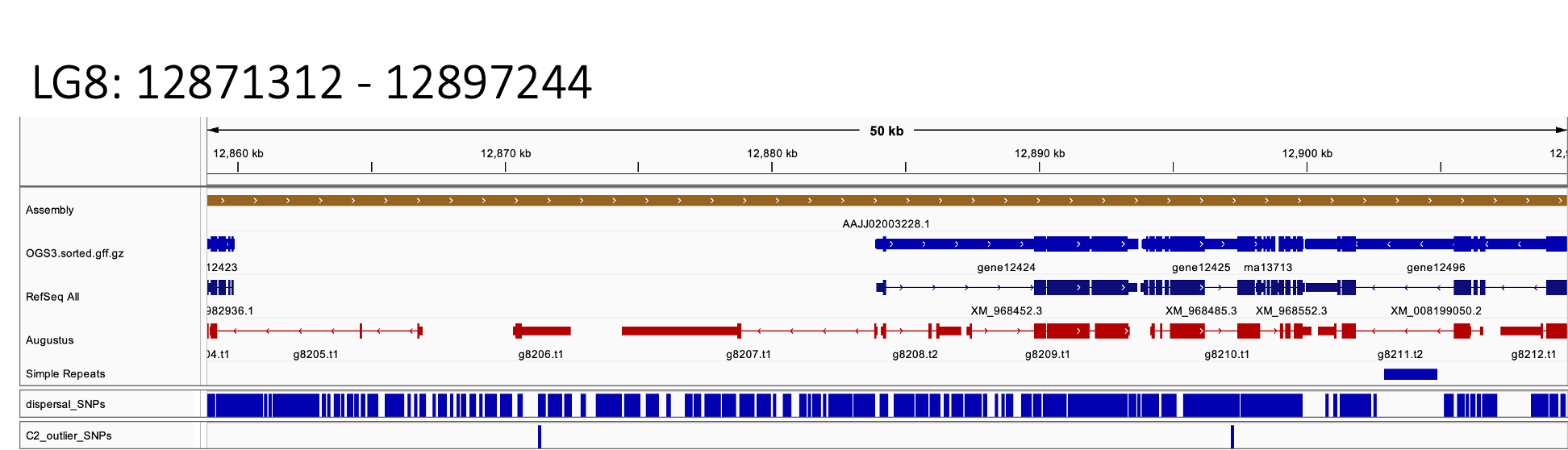

R)

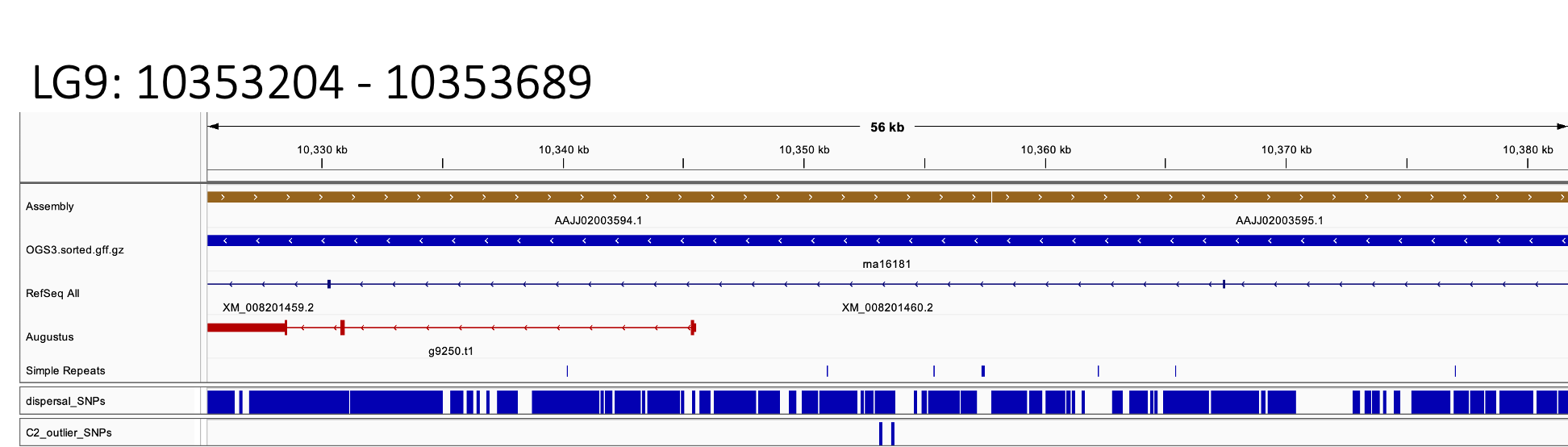

S)

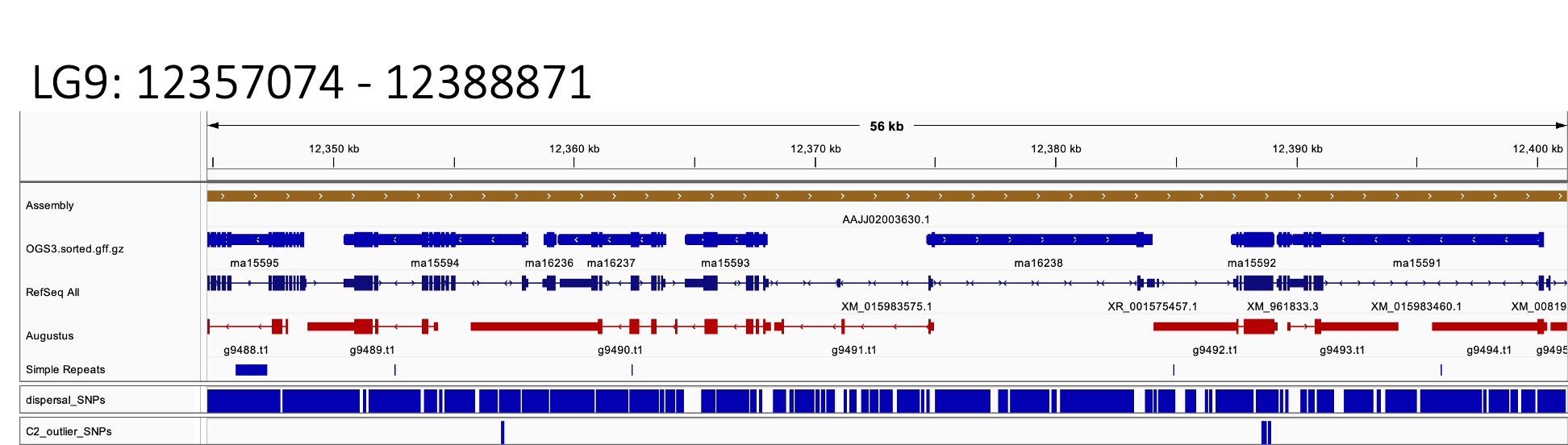

T)

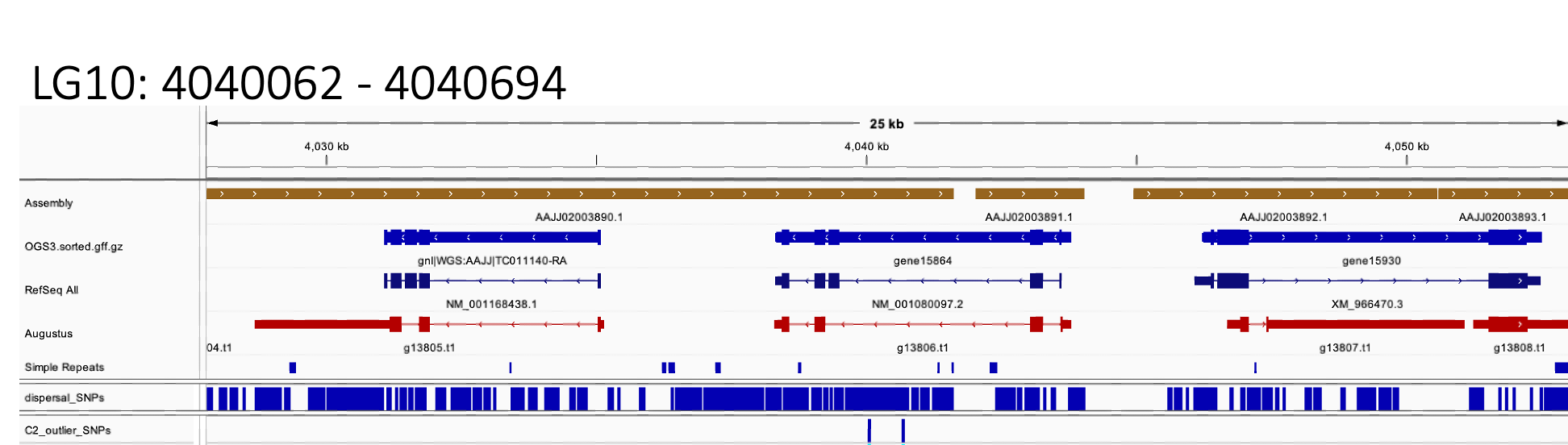

U)

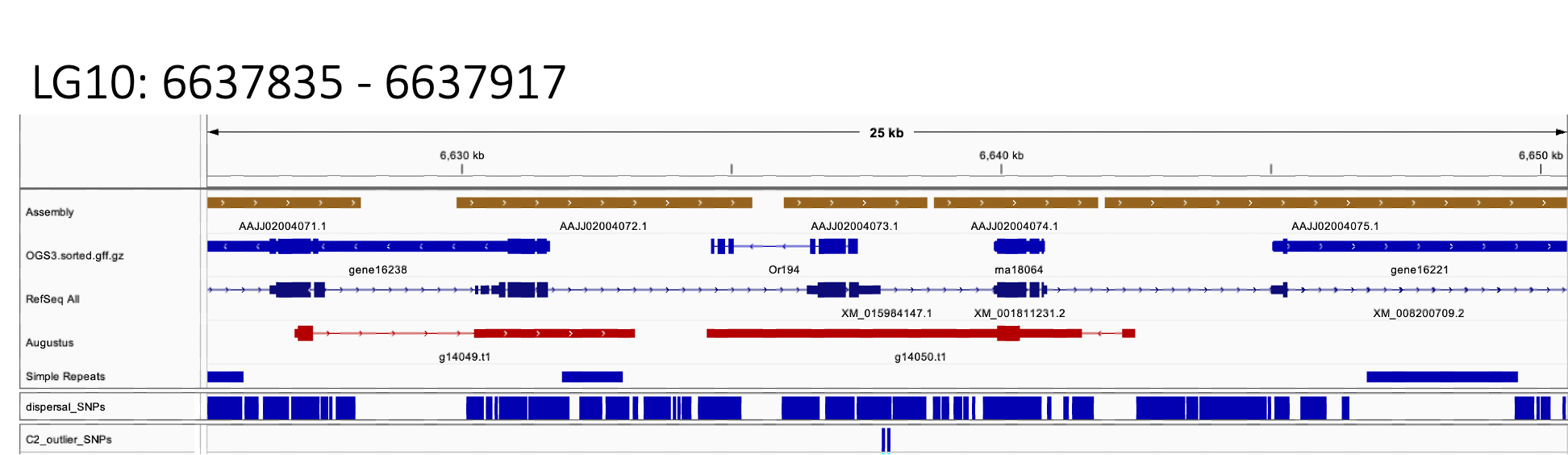

V)

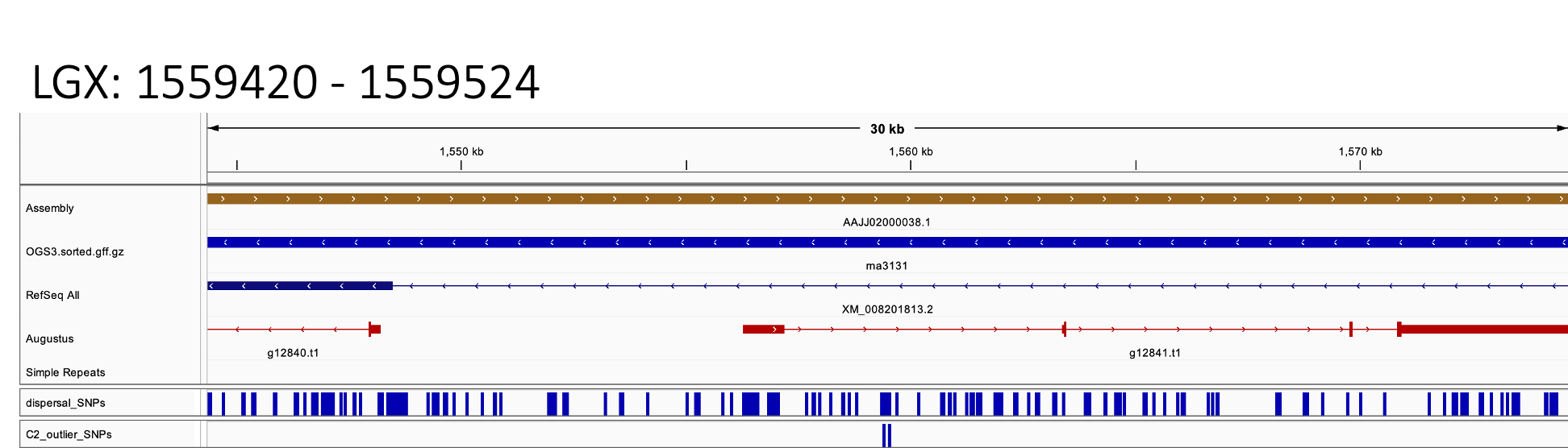

**Figure S5.** Each of 22 candidate regions identified by BayPass C2 outlier analysis, visualised with Integrative Genomics Viewer (Robinson *et al.* 2023). Panels A-V each show a single region, arranged by genomic position, and display, from top to bottom: assembled regions of the Tcas5.2 genome; OGS3 gene annotations; Ensembl RefSeqall gene annotations; Augustus gene annotations; the locations of all called SNPs across all populations in the dataset; and the locations of significant outlier SNPs from BayPass C2 outlier analysis.

**Table S2.** Genes identified as associated with dispersal behaviour in lines of *Tribolium castaneum* artificially selected for divergent dispersal propensity. All genes listed were found to be within candidate regions (C2), also indicated is whether each was within an outlier π window. Genes are ordered by the number of analyses supporting them, then by their genomic position. Information is provided from any Drosophila homologs listed in iBeetleBase and retrieved from FlyBase.

| **Genomic location (LG:POS)** | **Gene ID** | **C_2_** | **π** | **Flybase homolog ID** | **Flybase homolog name** | **Putative function** |
| --- | --- | --- | --- | --- | --- | --- |
| 2:9993610-10007042 | TC001033 | ✓ | ✓ | FBgn0043884 | Multiple ankyrin repeats single KH domain | Mediator of receptor tyrosine kinase (RTK) signaling, required for flight muscle sarcomere formation |
| 3:4347291-4365573 | TC003405 | ✓ | ✓ | FBgn0024315 | Picot | Predicted inorganic phosphate cotransporter |
| 3:4365581-4372318 | TC032364 | ✓ | ✓ | FBgn0004784 | Inactivation no afterpotential C | Encodes an eye-specific protein kinase C (PKC) involved in visual signaling; female gonad development; response to light |
| 3:5432673-5475367 | TC002977 | ✓ | ✓ | FBgn0035756 | Unc-13-4A | Predicted to be active in neurotransmitter secretory vesicle |
|  |  | ✓ | ✓ | FBgn0029727 | - | Transmembrane transporter activity; monoatomic anion transport |
| 4:4040766-4049514 | TC007470 | ✓ | ✓ | FBgn0264494 | - | Predicted to enable ATPase-coupled transmembrane transporter activity |
| 4:4052386-4056270 | TC008024 | ✓ | ✓ | FBgn0043841 | Virus-induced RNA 1 | Induced by viral infection, used as a marker of the induction of an antiviral response |
| 4:4056806-4058067 | TC030196 | ✓ | ✓ | - | - | - |
| 4:4059326-4060956 | TC030194 | ✓ | ✓ | - | - | - |
| 4:4060704-4061868 | TC030193 | ✓ | ✓ | - | - | - |
| 4:4063386-4066090 | TC032800 | ✓ | ✓ | - | - | - |
| 4:4063396-4064561 | TC030192 | ✓ | ✓ | - | - | - |
| 4:4067233-4071547 | TC032801 | ✓ | ✓ | FBgn0262738 | No receptor potential A | Phototransduction; photoentrainment; locomotor behaviour |
| 4:4071553-4072791 | TC030191 | ✓ | ✓ | - | - | - |
| 4:4073343-4074953 | TC008028 | ✓ | ✓ | FBgn0003965 | Vermilion | Heme-dependent dioxygenase, required during larval growth to control the level of potentially harmful free tryptophan; eye pigmentation |
| 4:4074968-4079671 | TC007467 | ✓ | ✓ | FBgn0031220 | - | Predicted to enable ATPase-coupled transmembrane transporter activity |
| 4:4082376-4091666 | TC007466 | ✓ | ✓ | FBgn0288229 | - | Enable ATPase-coupled transmembrane transporter activity. |
| 4:4091939-4098733 | TC008029 | ✓ | ✓ | FBgn0000463 | Delta | Regulates cell fate decisions and cell proliferation; leg/wing/antenna development; photoreceptor development; Notch signalling |
| 4:4098749-4100066 | TC007465 | ✓ | ✓ | FBgn0032727 | Betaine-homocysteine S-methyltransferase | Enable S-adenosylmethionine-homocysteine S-methyltransferase activity |
|  |  |  | ✓ | FBgn0032726 | - | Enable S-adenosylmethionine-homocysteine S-methyltransferase activity |
| 4:4100198-4101823 | TC008030 | ✓ | ✓ | FBgn0019957 | NADH dehydrogenase (ubiquinone) 42 kDa subunit | Encodes a subunit of complex I of the mitochondrial electron transport chain |
| 4:4103408-4103809 | TC008031 | ✓ | ✓ | FBgn0037579 | Cytochrome c oxidase subunit 7A-like | Involved in mitochondrial respirasome assembly and regulation of oxidative phosphorylation |
|  |  |  | ✓ | FBgn0085201 | Cytochrome c oxidase subunit 7A-like 2 | Involved in mitochondrial respirasome assembly and regulation of oxidative phosphorylation |
|  |  |  | ✓ | FBgn0040529 | Cytochrome c oxidase subunit 7A | Component of the cytochrome c oxidase, the last enzyme in the mitochondrial electron transport chain which drives oxidative phosphorylation |
| 4:4103878-4107194 | TC008032 | ✓ | ✓ | - | - | - |
| 6:1458748-1462133 | TC015494 | ✓ | ✓ | FBgn0023214 | ETS-domain lacking | Egfr signaling pathway regulation; sensory organ development; nuclear export |
| 6:1464348-1465586 | TC015211 | ✓ | ✓ | 33653 | - | Predicted to be involved in social behavior |
| 6:1466477-1467676 | TC015210 | ✓ | ✓ | 29851 | - | - |
| 6:1469091-1479796 | TC015209 | ✓ | ✓ | FBgn0034025 | Polypeptide N-Acetylgalactosaminyltransferase 1 | Catalyst of oligosaccharide biosynthesis |
| 6:1479915-1480926 | TC015208 | ✓ | ✓ | 29851 | - | - |
| 7:6750458-6783030 | TC033673 | ✓ | ✓ | FBgn0083946 | Lost boys | Key component of the nexin-dynein regulatory complex; sperm motility; sperm storage |
| 7:6785383-6787610 | TC016383 | ✓ | ✓ | 50401 | Distal antenna-young | Chromatin-binding protein required in spermatocytes for a normal gene expression profile |
|  |  |  | ✓ | 39283 | Distal antenna-related | Transcription factor with a role in the retinal determination (RD) network; eye/antenna development; nervous system formation |
| 7:6787398-6787895 | TC016382 | ✓ | ✓ | - | - | - |
| 7:6789568-6802399 | TC016381 | ✓ | ✓ | - | - | - |
| 7:6802465-6818299 | TC016380 | ✓ | ✓ | 44452 | Autophagy-related 2 | Encodes a protein known to be required for autophagy; wound healing |
| 7:6818541-6820106 | TC016342 | ✓ | ✓ | FBgn0010803 | Tryptophanyl-tRNA synthetase | Tryptophan-tRNA ligase activity; ATP binding. Dendrite morphogenesis |
| 8:12883884-12893699 | TC006687 | ✓ | ✓ | - | - | - |
| 8:12893824-12898338 | TC006688 | ✓ | ✓ | FBgn0040268 | Topoisomerase 3alpha | Encodes a type IA topoisomerase involved in junction dissolution during homologous recombination |
| 9:10278308-10422001 | TC034367 | ✓ | ✓ | FBgn0038881 | Argus | Autophagosome-lysosome fusion. mature autophagosome degradation. Loss of its autophagy function results in sleep loss |
| 9:12350445-12358135 | TC011872 | ✓ | ✓ | FBgn0250755 | - | Negative regulation of fatty acid biosynthetic process |
| 9:12358722-12359295 | TC034415 | ✓ | ✓ | - | - | - |
| 9:12359354-12363845 | TC034416 | ✓ | ✓ | 31213 | Galectin | Encodes a galactoside binding protein involved in synaptic target recognition |
|  |  |  | ✓ | 31214 | - | Predicted to enable carbohydrate binding activity and galactoside binding activity |
| 9:12364626-12368045 | TC011870 | ✓ | ✓ | 1297 | Kayak | Transcription factor involved in multiple biological processes; germ cell / eye development / response to wounding / locomotion / locomotor rhythm |
| 9:12374634-12384015 | TC034417 | ✓ | ✓ | - | - | - |
| 9:12374736-12383773 | TC034887 | ✓ | ✓ | - | - | - |
| 9:12387287-12389095 | TC011868 | ✓ | ✓ | - | - | - |
| 2:10190871-10206697 | TC032148 | ✓ | - | FBgn0266579 | Tau | Microtubule associated protein; human orthoglog implicated neuronal degradation |
| 2:10207381-10209696 | TC001051 | ✓ | - | FBgn0038666 | Smu1 spliceosomal factor | Encodes a spliceosomal protein required for normal neuromuscular junction development and function. |
| 2:10209941-10211960 | TC001052 | ✓ | - | FBgn0038047 | - | Predicted to enable DNA-binding transcription factor activity |
| 2:10215491-10217569 | TC000523 | ✓ | - | - | - | - |
| 2:10211972-10213759 | TC000525 | ✓ | - | - | - | - |
| 2:10213821-10215269 | TC000524 | ✓ | - | FBgn0028662 | Vacuolar H+ ATPase PPA1 subunit 1 | Encodes a protein involved in tracheal terminal branching |
| 2:10217617-10219009 | TC000522 | ✓ | - | FBgn0250732 | GST-containing FLYWCH zinc-finger protein | Predicted glutathione transferase activity |
| 2:13966054-13966404 | TC000259 | ✓ | - | - | - | - |
| 2:13969569-13970791 | TC001287 | ✓ | - | FBgn0035980 | Mitochondrial ribosome recycling factor 1 | Predicted to enable ribosomal large subunit binding activity |
| 2:13970858-13973137 | TC000258 | ✓ | - | FBgn0032492 | Proteasome alpha6 subunit, Testis-specific | Encodes a testes-specific subunit of the 26S proteasome; spermatogenesis |
|  |  |  | - | FBgn0250843 | Proteasome alpha6 subunit | Proteasome has an ATP-dependent proteolytic activity' adult lifespan |
| 2:13974218-13974362 | TC001288 | ✓ | - | FBgn0027660 | Bloated tubules | Encodes a member of the neurotransmitter symporter family |
| 2:13976786-13985962 | TC034566 | ✓ | - | FBgn0036274 | - | Enable DNA-binding transcription factor; neuron differentiation; regulation of transcription |
| 2:13994799-13995468 | TC000256 | ✓ | - | - | - | - |
| 2:14001779-14005903 | TC001290 | ✓ | - | - | - | - |
| 2:14008656-14015259 | TC001291 | ✓ | - | FBgn0052105 | LIM homeobox transcription factor 1 alpha | Enable DNA-binding transcription factor; neuron differentiation; regulation of transcription |
| 2:14015621-14016244 | TC000255 | ✓ | - | FBgn0036277 | - | RNA binding activity; regulation of alternative mRNA splicing |
| 2:14016336-14019511 | TC001292 | ✓ | - | FBgn0041147 | Imaginal discs arrested | Encodes ubiquitin ligase that regulates mitotic metaphase/anaphase transition |
| 2:14023666-14033821 | TC001293 | ✓ | - | FBgn0031407 | - | - |
|  |  | ✓ | - | FBgn0054049 | - | - |
|  |  | ✓ | - | FBgn0031412 | - | Involved in sexual reproduction |
|  |  | ✓ | - | FBgn0051482 | - | - |
|  |  | ✓ | - | FBgn0051286 | - | - |
| 2:14033838-14073163 | TC032248 | ✓ | - | FBgn0261698 | Slowpoke 2 | Encodes a channel involved in potassium ion transmembrane transport |
|  |  |  | - | FBgn0262514 | Vacuolar H+ ATPase PPA1 subunit 2 | Predicted to contribute to proton-transporting ATPase activity, rotational mechanism |
| 4:5843998-5854519 | TC008154 | ✓ | - | FBgn0001404 | Egghead | Glycosphingolipid biosynthesis; germ cell development |
| 4:5855090-5856776 | TC007345 | ✓ | - | - | - | - |
| 4:5859049-5859492 | TC008155 | ✓ | - | - | - | - |
| 4:5859118-5871357 | TC008156 | ✓ | - | FBgn0259146 | Fire dancer | Encodes a protein involved in heat resistance |
| 4:5871397-5876636 | TC007344 | ✓ | - | FBgn0026015 | Topoisomerase 3β | DNA binding; transcription and translation regulation |
| 4:5876728-5879344 | TC008157 | ✓ | - | FBgn0003520 | Staufen | Encodes a double-stranded RNA binding protein involved in mRNA localization; long-term memory; oogenesis |
| 4:5883188-5887835 | TC032826 | ✓ | - | - | - | - |
| 4:5895151-5895408 | TC008159 | ✓ | - | - | - | - |
| 5:1468679-1472342 | TC010933 | ✓ | - | FBgn0250815 | Jonah 65Aiv | Enables serine hydrolase activity; innate immune response |
|  |  |  | - | FBgn0036264 |  | Serine-type endopeptidase activity; innate immune response |
|  |  |  | - | FBgn0052383 | Sphinx1 | Serine protease, strongly expressed in the male accessory gland; sexual reproduction |
|  |  |  | - | FBgn0052382 | Sphinx2 | Serine protease, strongly expressed in the male accessory gland; sexual reproduction |
| 5:1477499-1478648 | TC010934 | ✓ | - | FBgn0250815 | Jonah 65Aiv | Enables serine hydrolase activity; innate immune response |
|  |  |  | - | FBgn0036264 |  | Serine-type endopeptidase activity; innate immune response |
|  |  |  | - | FBgn0052383 | Sphinx1 | Serine protease, strongly expressed in the male accessory gland; sexual reproduction |
|  |  |  | - | FBgn0052382 | Sphinx2 | Serine protease, strongly expressed in the male accessory gland; sexual reproduction |
| 5:1481411-1486042 | TC010910 | ✓ | - | FBgn0250815 | Jonah 65Aiv | Enables serine hydrolase activity; innate immune response |
|  |  |  | - | FBgn0036264 |  | Serine-type endopeptidase activity; innate immune response |
|  |  |  | - | FBgn0052383 | Sphinx1 | Serine protease, strongly expressed in the male accessory gland; sexual reproduction |
|  |  |  | - | FBgn0052382 | Sphinx2 | Serine protease, strongly expressed in the male accessory gland; sexual reproduction |
| 5:1488476-1489657 | TC010935 | ✓ | - | 250815 | Jonah 65Aiv | Enables serine hydrolase activity; innate immune response |
|  |  |  | - | 36264 |  | Serine-type endopeptidase activity; innate immune response |
|  |  |  | - | 52383 | Sphinx1 | Serine protease, strongly expressed in the male accessory gland; sexual reproduction |
|  |  |  | - | 52382 | Sphinx2 | Serine protease, strongly expressed in the male accessory gland; sexual reproduction |
| 5:1492716-1493874 | TC010908 | ✓ | - | 250815 | Jonah 65Aiv | Enables serine hydrolase activity; innate immune response |
|  |  |  | - | 36264 |  | Serine-type endopeptidase activity; innate immune response |
|  |  |  | - | 52383 | Sphinx1 | Serine protease, strongly expressed in the male accessory gland; sexual reproduction |
|  |  |  | - | 52382 | Sphinx2 | Serine protease, strongly expressed in the male accessory gland; sexual reproduction |
| 5:1496434-1497448 | TC033014 | ✓ | - | - | - | - |
| 5:1500054-1504630 | TC010936 | ✓ | - | 250815 | Jonah 65Aiv | Enables serine hydrolase activity; innate immune response |
|  |  |  | - | 36264 |  | Serine-type endopeptidase activity; innate immune response |
|  |  |  | - | 52383 | Sphinx1 | Serine protease, strongly expressed in the male accessory gland; sexual reproduction |
|  |  |  | - | 52382 | Sphinx2 | Serine protease, strongly expressed in the male accessory gland; sexual reproduction |
| 5:1503565-1504914 | TC033013 | ✓ | - | - | - | - |
| 5:1508306-1509471 | TC010937 | ✓ | - | - | - | - |
| 5:1512846-1515333 | TC010938 | ✓ | - | - | - | - |
| 5:1521986-1527854 | TC033012 | ✓ | - | 250815 | Jonah 65Aiv | Enables serine hydrolase activity; innate immune response |
|  |  |  | - | 36264 |  | Serine-type endopeptidase activity; innate immune response |
|  |  |  | - | 52383 | Sphinx1 | Serine protease, strongly expressed in the male accessory gland; sexual reproduction |
|  |  |  | - | 52382 | Sphinx2 | Serine protease, strongly expressed in the male accessory gland; sexual reproduction |
| 5:1528136-1529203 | TC010939 | ✓ | - | 250815 | Jonah 65Aiv | Enables serine hydrolase activity; innate immune response |
|  |  |  | - | 36264 |  | Serine-type endopeptidase activity; innate immune response |
|  |  |  | - | 52383 | Sphinx1 | Serine protease, strongly expressed in the male accessory gland; sexual reproduction |
|  |  |  | - | 52382 | Sphinx2 | Serine protease, strongly expressed in the male accessory gland; sexual reproduction |
| 5:1531166-1532316 | TC010940 | ✓ | - | - | Serine protease P121 | May be involved in wound healing |
| 5:1535151-1542504 | TC033011 | ✓ | - | FBgn0250815 | Jonah 65Aiv | Enables serine hydrolase activity; innate immune response |
|  |  |  | - | FBgn0036264 |  | Serine-type endopeptidase activity; innate immune response |
|  |  |  | - | FBgn0052383 | Sphinx1 | Serine protease, strongly expressed in the male accessory gland; sexual reproduction |
|  |  |  | - | FBgn0052382 | Sphinx2 | Serine protease, strongly expressed in the male accessory gland; sexual reproduction |
| 5:1710318-1732029 | TC010898 | ✓ | - | FBgn0261258 | Regeneration | Carbohydrate binding; tissue regeneration |
| 5:3791386-3796339 | TC013804 | ✓ | - | FBgn0000120 | Arrestin 1 | Photoreceptor maintenance and smell perception |
| 5:3799189-3808589 | TC013803 | ✓ | - | 10583 | Dreadlocks | Adapter protein inking cell surface receptor tyrosine phosphorylation to downstream signaling pathways and effectors; protein binding; insulin receptor signalling |
| 6:9247748-9256186 | TC014879 | ✓ | - | FBgn0012051 | Calpain-A | Encodes a calcium-dependent modulatory protease; adult lifespan; muscle development; larval locomotion |
|  |  |  | - | FBgn0025866 | Calpain-B | Calcium-regulated non-lysosomal thiol-protease |
|  |  |  | - | FBgn0260450 | Calpain-C | Encodes a calcium-dependent cysteine protease |
| 6:9259798-9260781 | TC015841 | ✓ | - | 39203 | Juvenile hormone binding protein 12 | - |
|  |  |  | - | 38850 | Juvenile hormone binding protein 15 | - |
| 6:9266428-9267445 | TC015842 | ✓ | - | 39203 | Juvenile hormone binding protein 12 | - |
| 6:9270702-9271646 | TC014878 | ✓ | - | 39203 | Juvenile hormone binding protein 12 | - |
| 6:9275948-9276894 | TC015843 | ✓ | - | 39203 | Juvenile hormone binding protein 12 | - |
| 6:9279888-9281209 | TC015844 | ✓ | - | 39203 | Juvenile hormone binding protein 12 | - |
| 6:9286120-9305396 | TC014877 | ✓ | - | FBgn0034909 | Pippin | Transmembrane transport of sugars |
| 7:2895583-2925437 | TC033574 | ✓ | - | FBgn0085390 | Diacyl glycerol kinase | ATP-dependent diacylglycerol kinase activity; calcium ion binding. May be involved in muscle function and regulating neuron signalling |
| 7:8224581-8233768 | TC009038 | ✓ | - | FBgn0086784 | Stambha A | Synaptic vesicle endocytosis; phototransduction; and synaptic vesicle exocytosis |
| 8:9611455-9615202 | TC015261 | ✓ | - | FBgn0003510 | Serendipity α | cellularization of the syncytial blastoderm embryo |
|  |  |  | - | FBgn0033348 | Spitting Image | Encodes a stabilizing protein in the early embryo |
| 8:9615206-9627710 | TC030897 | ✓ | - | - | - | - |
| 8:9623376-9626906 | TC015262 | ✓ | - | - | - | - |
| 8:9627795-9629240 | TC015264 | ✓ | - | FBgn0036856 | Methyltransferase like 5 | Methylates the 6th position of adenine in 18S rRNA |
| 8:9627856-9628159 | TC015263 | ✓ | - | 259720 | - | Predicted to be involved in DNA repair |
| 8:9630888-9636133 | TC033340 | ✓ | - | FBgn0000594 | Esterase P | Enables carboxylesterase activity |
|  |  |  | - | FBgn0000592 | Esterase 6 | Carboxylesterase; odorant sensing; pheromone response; courtship behaviour, egg-laying behaviour, sexual reproduction |
| 8:9636275-9640528 | TC033339 | ✓ | - | FBgn0000594 | Esterase P | Enables carboxylesterase activity |
|  |  |  | - | FBgn0000592 | Esterase 6 | Carboxylesterase; odorant sensing; pheromone response; courtship behaviour, egg-laying behaviour, sexual reproduction |
| 10:4038317-4043813 | TC011139 | ✓ | - | FBgn0027600 | Obstructor-B | Chitin binding activity |
| X:1533167-1577906 | TC004127 | ✓ | - | - | - | Involved in nervous system development |

Ahrens CW, Rymer PD, Stow A, Bragg J, Dillon S, Umbers KDL, *et al.* (2018). The search for loci under selection: trends, biases and progress. *Mol Ecol* **27**: 1342–1356.

Broad Institute (2019). “Picard Toolkit.” 2019. GitHub Repository. <https://broadinstitute.github.io/picard/>

Gautier M (2015). Genome-Wide Scan for Adaptive Divergence and Association with Population-Specific Covariates. *Genetics* **201**: 1555–1579.

Lotterhos KE (2019). The Effect of Neutral Recombination Variation on Genome Scans for Selection. *G3*  **9**: 1851–1867.

Quinlan AR, Hall IM (2010). BEDTools: a flexible suite of utilities for comparing genomic features. *Bioinformatics* **26**: 841–842.

Robinson JT, Thorvaldsdottir H, Turner D, Mesirov JP (2023). igv.js: an embeddable JavaScript implementation of the Integrative Genomics Viewer (IGV). *Bioinformatics* **39**: btac830.
